# Comprehensive Assessment of Blood-Brain Barrier Opening and Sterile Inflammatory Response: Unraveling the Therapeutic Window

**DOI:** 10.1101/2023.10.23.563613

**Authors:** Payton Martinez, Jane J. Song, Francis G. Garay, Kang-Ho Song, Toni Mufford, Jenna Steiner, John DeSisto, Nicholas Ellens, Natalie J. Serkova, Adam L. Green, Mark Borden

## Abstract

Microbubbles (MBs) combined with focused ultrasound (FUS) have emerged as a promising noninvasive technique to permeabilize the blood-brain barrier (BBB) for drug delivery to the brain. However, the safety and biological consequences of BBB opening remain incompletely understood. This study investigates the effects of varying microbubble volume doses (MVD) and ultrasound mechanical indices (MI) on BBB opening and the sterile inflammatory response (SIR) using high-resolution ultra-high field MRI-guided FUS in mouse brains. The results demonstrate that both MVD and MI significantly influence the extent of BBB opening, with higher doses and mechanical indices leading to increased permeability. Moreover, RNA sequencing reveals upregulated inflammatory pathways and immune cell infiltration after BBB opening, suggesting the presence and extent of SIR. Gene set enrichment analysis identifies 12 gene sets associated with inflammatory responses that are upregulated at higher doses of MVD or MI. A therapeutic window is established between significant BBB opening and the onset of SIR, providing operating regimes for avoiding each three classes of increasing damage from stimulation of the NFκB pathway via TNFL signaling to apoptosis. This study contributes to the optimization and standardization of BBB opening parameters for safe and effective drug delivery to the brain and sheds light on the underlying molecular mechanisms of the sterile inflammatory response.

**Significance Statement:** The significance of this study lies in its comprehensive investigation of microbubble-facilitated focused ultrasound for blood-brain barrier (BBB) opening. By systematically exploring various combinations of microbubble volume doses and ultrasound mechanical indices, the study reveals their direct impact on the extent of BBB permeability and the induction of sterile inflammatory response (SIR). The establishment of a therapeutic window between significant BBB opening and the onset of SIR provides critical insights for safe and targeted drug delivery to the brain. These findings advance our understanding of the biological consequences of BBB opening and contribute to optimizing parameters for clinical applications, thus minimizing potential health risks, and maximizing the therapeutic potential of this technique.

## Introduction

Since the discovery of noninvasively permeabilizing the blood-brain barrier (BBB) through microbubbles and focused ultrasound (MB+FUS) (1), multiple efforts have been undertaken to shuttle drugs such as chemotherapeutics, antibodies, and other therapeutic molecules and carriers into the brain parenchyma (2, 3). This has been reflected in the growing number of clinical trials registered per year (4). With the growing number of applications MB+FUS has been employed for, most attention has been devoted to optimizing blood-brain barrier opening (BBBO) parameters on the ultrasound side, while conventional echocardiography contrast agent microbubbles with polydisperse size distributions have been employed at or near their clinical dose. As the field moves to more sophisticated microbubble formulations with longer circulation persistence and monodisperse size distributions, there is a need to elucidate the effects of microbubble dose on BBBO. Additionally, the full extent of the sterile inflammatory response (SIR) from BBBO still needs to be characterized, in terms of both ultrasound and microbubble doses. Considering this, the safety of BBBO in the future needs to be considered to minimize health concerns and mitigate harmful secondary effects (5).

The BBB is crucial in maintaining homeostasis and is our first line of defense against pathogens and noxious insults, which would cause considerable damage if permitted to cross into the brain parenchyma (6). To avoid the incidental passage of undesirable molecules, the BBB is made up of a basal layer of endothelial cells, which selectively exclude hydrophilic molecules larger than 400 Daltons (7, 8),. Tight junction proteins such as occludins, claudins and junctional adhesion molecules exist on endothelial cell membranes and form complexes to fasten together neighboring cells (9, 10). Additionally, pericytes and astrocytic end feet processes envelop the vasculature, regulating vascular constriction and ensuring proper maintenance of the barrier (11–13). Collectively, these constituents form a structure referred to as the neurovascular unit.

Microbubbles (MBs) are ultrasound (US) responsive colloidal particles that have a gas core encapsulated in a phospholipid monolayer shell (14–17). These 1-10 µm diameter spheres experience an isotropic but dynamic pressure field as the ultrasound wave (∼1 mm wavelength) passes over, resulting in volumetric oscillations at MHz frequency within the ultrasound focal region (16). The ultrasound is typically fixed to lower frequency (F, ∼1 MHz for mice and ∼0.2-0.5 MHz for humans) to ensure transcranial propagation, but the amplitude (PNP, peak negative pressure) can be varied to produce a range of bioeffects. The dose of ultrasound can thus be adjusted to achieve the desired mechanical index (MI = PNP/F^1/2^) (18). With increasing MI, the microbubble acoustic behavior progresses from mild harmonic oscillations to violent inertial implosions (19–21). It is generally considered safe to avoid inertial oscillations (22, 23). In the context of BBBO, harmonic MB oscillations were found to pry apart tight junction proteins, creating transient pores in the brain endothelium and allowing blood-borne molecules to extravasate,(24–27). Additional ultrasound parameters that can be optimized include pulse repetition frequency, pulse length and total sonication time, as well as details of the ultrasound beam and focal region (28, 29).

As the microbubble is the acoustic actuator that captures the acoustic energy and uses it to produce localized mechanical work on the endothelium, it is also an important parameter that must considered. Of particular interest is the size and concentration, which can be quantified as the injected microbubble volume dose (MVD, µL/kg) (30, 31). When matching MVD, microbubbles of different sizes were found to produce similar pharmacokinetic profiles (32, 33), acoustic response as measured by passive cavitation detection (34), and extent of BBB opening (30). Thus, MVD and MI serve as relevant microbubble and ultrasound dosing parameters, forming two axes from which a window of safety and efficacy can be discerned.

Despite rigorous technical characterization over the years, one of the most understudied areas is related to the biological consequences of BBBO. On a larger scale, it has been observed that temporarily permeabilizing the BBB can induce microhemorrhages, transient edema and even cell death (4). Upon closer examination, transcriptomic analyses of the parenchymal microenvironment post MB+FUS have revealed upregulation of several major inflammatory pathways, mostly notably the NFκB pathway (28, 35). While occurring in the absence of an active infection, this event has been labeled as the sterile inflammatory response (SIR) and is initiated when damage-associated molecular patterns are released from injured cells (36–38). These include ATP, uric acid, DNA and HMBG proteins, which bind to pattern recognition receptors and provoke the immune system (39, 40). Subsequently, proinflammatory cytokines such as TNFL and IL2 are released from inflammasomes and stimulate the activation of innate immunity (41). Several studies have implicated the hallmark activation of the NFκB pathway in the persistence of SIR; however, the direct mechanism of activation and associated pathways remain unclear (42–44). Moreover, microglia, the primary immune cells of the brain, migrate to the source of inflammation and release cytokines, signaling the recruitment of peripheral immune cells and other cell types in the area (35, 45). Peripheral immune cells such as CD68+ macrophages circulate in the meninges and lymphatic system via chemotaxis, migrate to the area to investigate and resolve the inflammatory response (35). Despite BBB closure within a 24-hour window, peripheral immune cells have been known to continue extravasating past the BBB (46, 47).

Here, we report on a study to determine the extent of BBBO and SIR using a combination of MI doses (0.2 - 0.6 MPa/MHz^1/2^) and MVD (0.1 – 40 µL/kg). By using MRI-guided FUS, we sonicated the right mouse striatum and collected brain samples for bulk RNA sequencing 6 hours post-sonication. We defined BBBO as 15% increase in signal intensity after gadolinium bolus injection over a 1 mm^3^ volume in contrast enhanced T1-weighted MRI (CE-T1w MRI). Additionally, we defined SIR according to three classes defined by normalized enrichment scores (NES > 1.65) (Class I: TNFL signaling via NFκB gene set; Class II: both the TNFL signaling via NFκB and Inflammatory response gene sets; Class III: TNFL signaling via NFκB, Inflammatory response, and a damage-associated marker, apoptosis). Using these criteria, we developed a therapeutic window of ultrasound MI and microbubble MVD between the onset of BBBO and the onset of SIR, for each of the three classes. These windows will help to determine safe and efficacious MB+FUS parameters for BBBO in various applications.

## Results

### Microbubbles are Monodisperse in Size

Microbubbles were isolated to a uniform size of 3-µm diameter, as seen under brightfield microscopy (Figure 1A). The size distribution of the microbubbles was found to be monodisperse, with narrow peaks observed in both the number- and volume-weighted distributions (Figure 1B). The mean diameters for the number- and volume-weighted distributions were 3.3 µm and 3.7 µm, respectively. The 10^th^ and 90^th^ percentiles in diameter were determined to be 2.53 µm and 4.19 µm respectively (Table S1). To ensure consistent microbubble volume dose injection, each microbubble batch was analyzed. Figure 1C illustrates the relationship between microbubble concentration and volume, which was integrated to calculate the gas volume fraction (LMB). For a basis concentration of 10^10^ MBs/mL, the mean LMB was determined to be 17 µL/mL (Table S1). MVD is calculated by multiplying LMB by the fluid volume dose (mL/kg) injected intravenously into the subject.

**Figure 1.**
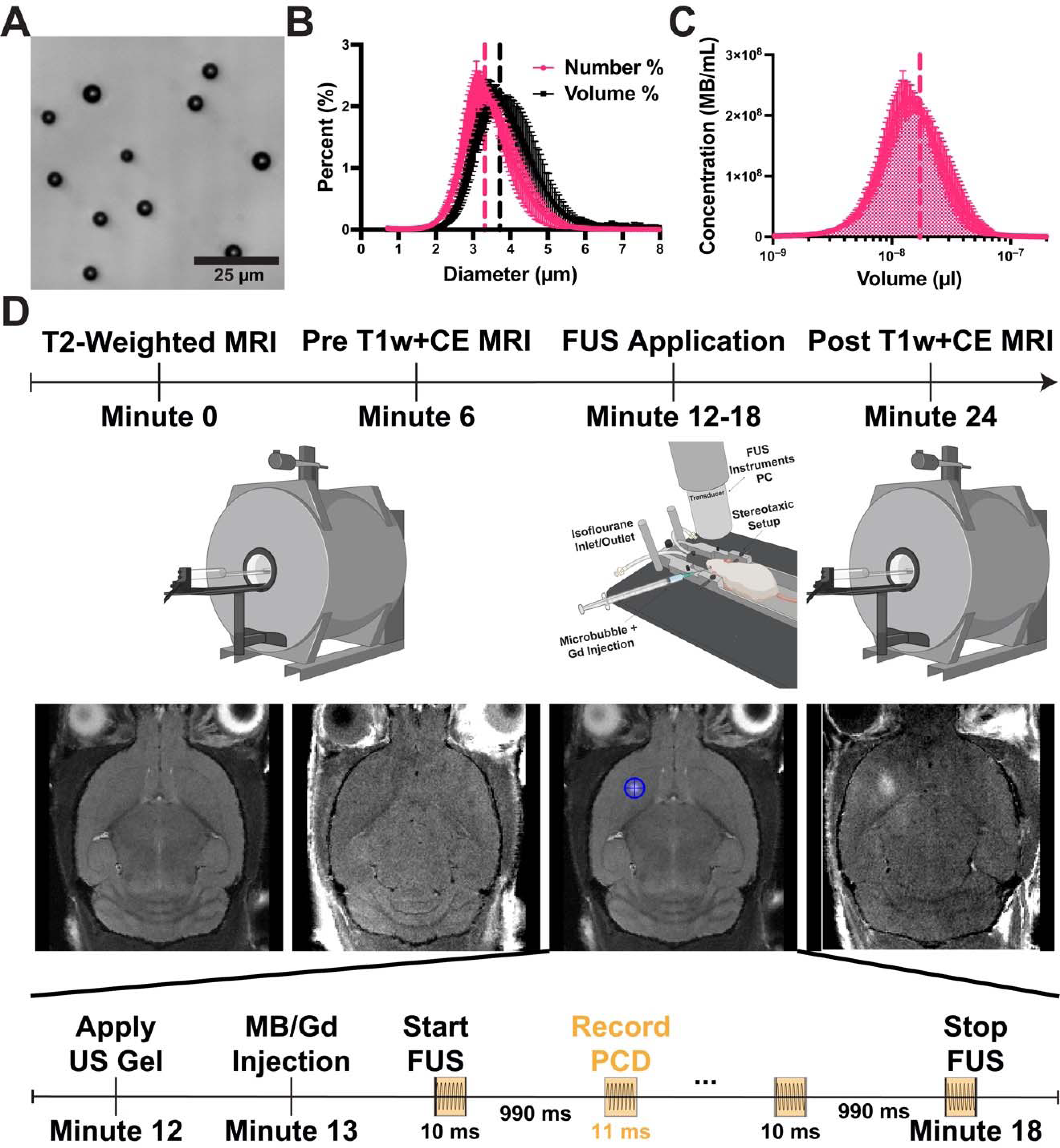
Microbubble Characterization and Treatment Timeline. (A) Brightfield image of isolated microbubbles (3 µm) in size. (B) Number- and volume-weighted size distributions. Vertical dotted lines represent mean values. (C) Microbubble concentration plotted against volume at a basis concentration of 10^10^ MBs/mL. The shaded region under the curve represents the gas volume fraction. (D) Illustration of the MB+FUS treatment timeline. Initially, mice were imaged using T2w and CE-T1w MRI. Subsequently, they were moved to the RK-50 system to receive MB+FUS treatment. A more detailed timeline is provided at the bottom of panel D. After treatment, mice were imaged again with CE-T1w MRI. Data are presented as mean ± standard deviation (*n* = 5).

### Blood-Brain Barrier Opening is Dependent on Both MVD and MI

Before the FUS treatment, all mice underwent T1w MRI after an injection of MultiHance (gadolinium contrast, CE-T1w MRI) and T2-weighted (T2w) MRI to confirm the integrity of the BBB and normal morphology of the brain, respectively. The CE-T1wMRI revealed an intact BBB prior to FUS procedure, as seen of minimal to no enhancement after MultiHance injection; quantification of BBBO was performed on post-FUS CE-T1w MRI, based on the changes in T1 signal intensities in T1w vs CE-T1w MRI as illustrated in Figure 2A. The contrast enhancement difference between the contralateral hemispheres before focused ultrasound treatment was determined to be 0.05 ± 6% (mean ± standard deviation) (Supplemental Figure 1). T2w MRI confirmed normal brain morphology in all animals.

**Figure 2.**
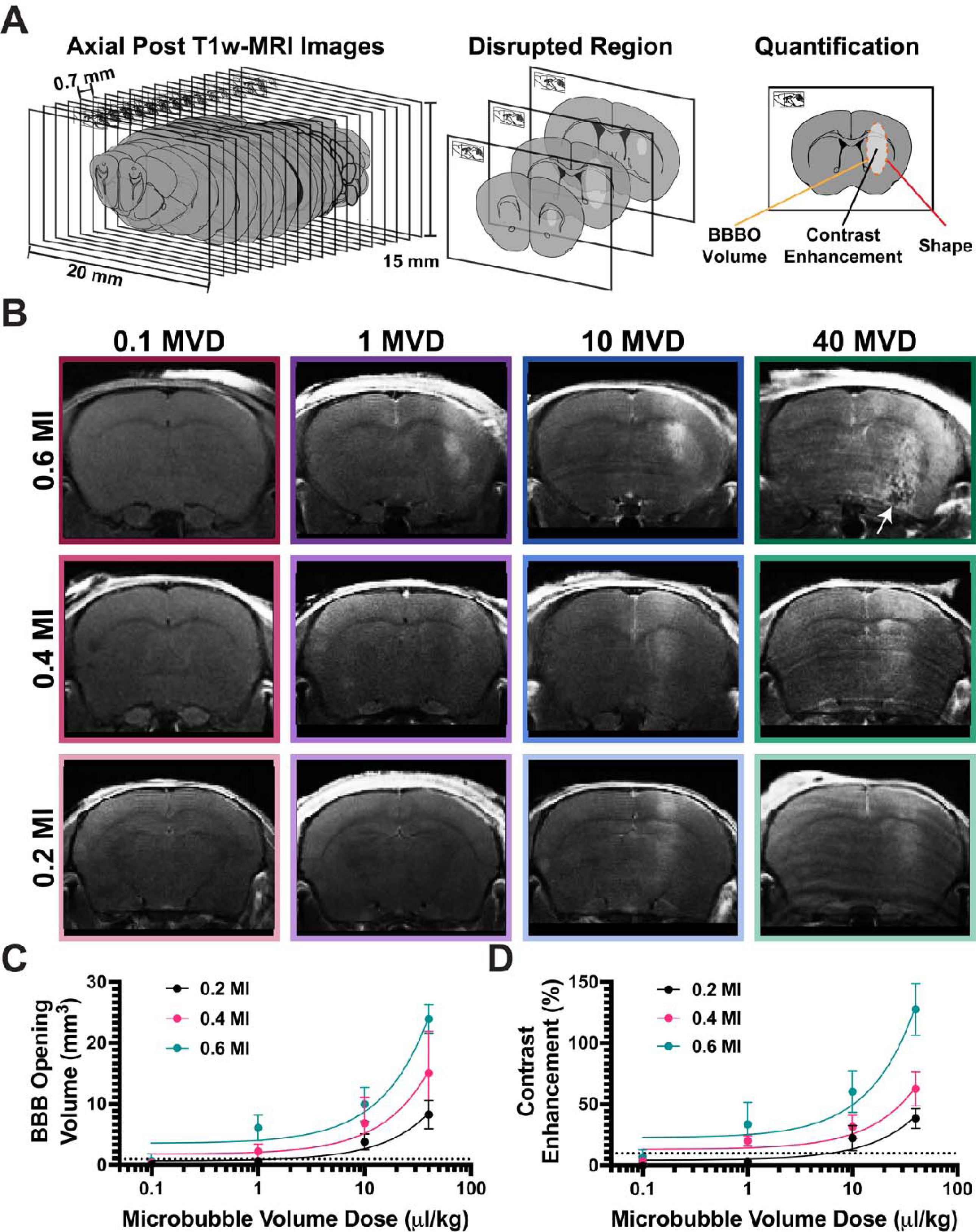
Assessment of BBBO using Gadolinium Contrast Enhancement. (A) Cartoon depicting post-CE-T1w MRI image analysis. (B) Representative images of post-FUS MRI for each MI/MVD dose, captured at the peak BBBO area. (C) Quantification of BBBO volume (*n* = 3). Linear regression was performed for each mechanical index, resulting in R squared values of 0.86, 0.71, and 0.91 for 0.2, 0.4, and 0.6 MI, respectively. The X-axis is presented on a logarithmic scale. (D) Quantification of BBBO contrast enhancement (*n* = 3). Linear regression was conducted for each mechanical index, resulting in R squared values of 0.79, 0.82, and 0.87 for 0.2, 0.4, and 0.6 MI, respectively. The X-axis is presented on a logarithmic scale. Data are presented as mean ± standard deviation.

Figure 2B shows representative images of BBBO for all twelve doses (MI/MVD). Within these experimental parameters, a clear onset of BBBO occurs at MVDs greater than 1 µL/kg at 0.2 MI, and MVDs greater than 0.1 µL/kg at both 0.4 and 0.6 MI. Irregular morphology was observed at the highest dose, as indicated by loss of brain structure in hypointense areas in T1w MRI (white arrow in Figure 2B). As the MI or MVD was increased, both the BBBO volume and amplitude increase, as seen by increased hyperintense areas and the relative signal intensity increase (Figures 2C and D). Linear trends between brain contrast enhancement in BBBO (as referred to both the enhanced volume and the relative signal intensity increase) and the MI/MVD were observed, yielding significant (p < 0.05) trends for all MIs (Figure 2C and D). Transposed plots comparing volume and contrast enhancement to MI are shown in Supplemental Fig. 2A and B. Multiple regression analysis (MI+MVD) between BBBO contrast enhancement resulted in R^2^ = 0.86, although R^2^ increased to 0.90 when analyzed as MI*MVD. This was similar for BBBO volume, with R^2^ moving from 0.89 (MI+MVD) to 0.94 (MI*MVD). Using FIJI (NIH, Bethesda, Maryland), a round and circle score was given to each shape of BBBO, and no significant differences were found between any dose pair (Supplemental Fig. S3). Based on the significance testing of pre-FUS controls (0.05 ± 6% (mean ± standard deviation), Supplemental Fig. 1), significant BBBO after FUS was defined as a 15% contrast enhancement over a 1 mm^3^ volume. These intensity and volume thresholds are represented as dotted lines in Figures 2C and D.

### Immunohistochemistry Shows Increased BBBO Leads to Stronger Immune Cell Infiltration

In addition to RNA sequencing, a subset of mice underwent immunohistochemistry (IHC) to track immune cell infiltration at the site of MB+FUS treatment. Five main markers for immune cells were utilized, including GFAP, Iba1, CD68, CD4 and CD8 staining, astrocytes, microglia, peripheral macrophages, helper T-cells and cytotoxic T-cells. As a major effector protein, NFκB was used as a marker for detecting inflammatory hotspots in the region of sonication. Representative close-up images of each marker, along with a DAPI nuclear stain, are shown in Figure 3A at the respective resolution for each analyzed image. Histology was conducted at the three MIs used in the study (0.2, 0.4 and 0.6) and an MVD of 10 µL/kg. The images indicate infiltration of immune cells as the MI increases (Figure 3B). MRI images showing the location of BBBO through increased MRI contrast correlate with the increased fluorescence of the IHC markers at these locations. Notably, at the highest MI (0.6), there is evident migration of astrocytes and microglia to the injured region. Hematoxylin and eosin (H&E) staining was also performed at each MI (Figure 3C). Apart from small amounts of red blood cell extravasation at the highest dose (black arrows), there was little difference observed in the morphology. Additionally, CD44 (activated immune cells) showed increased expression with increasing MI, while Luxol fast blue (myelin integrity) showed no differences between the groups (Supplemental Figure 4).

**Figure 3.**
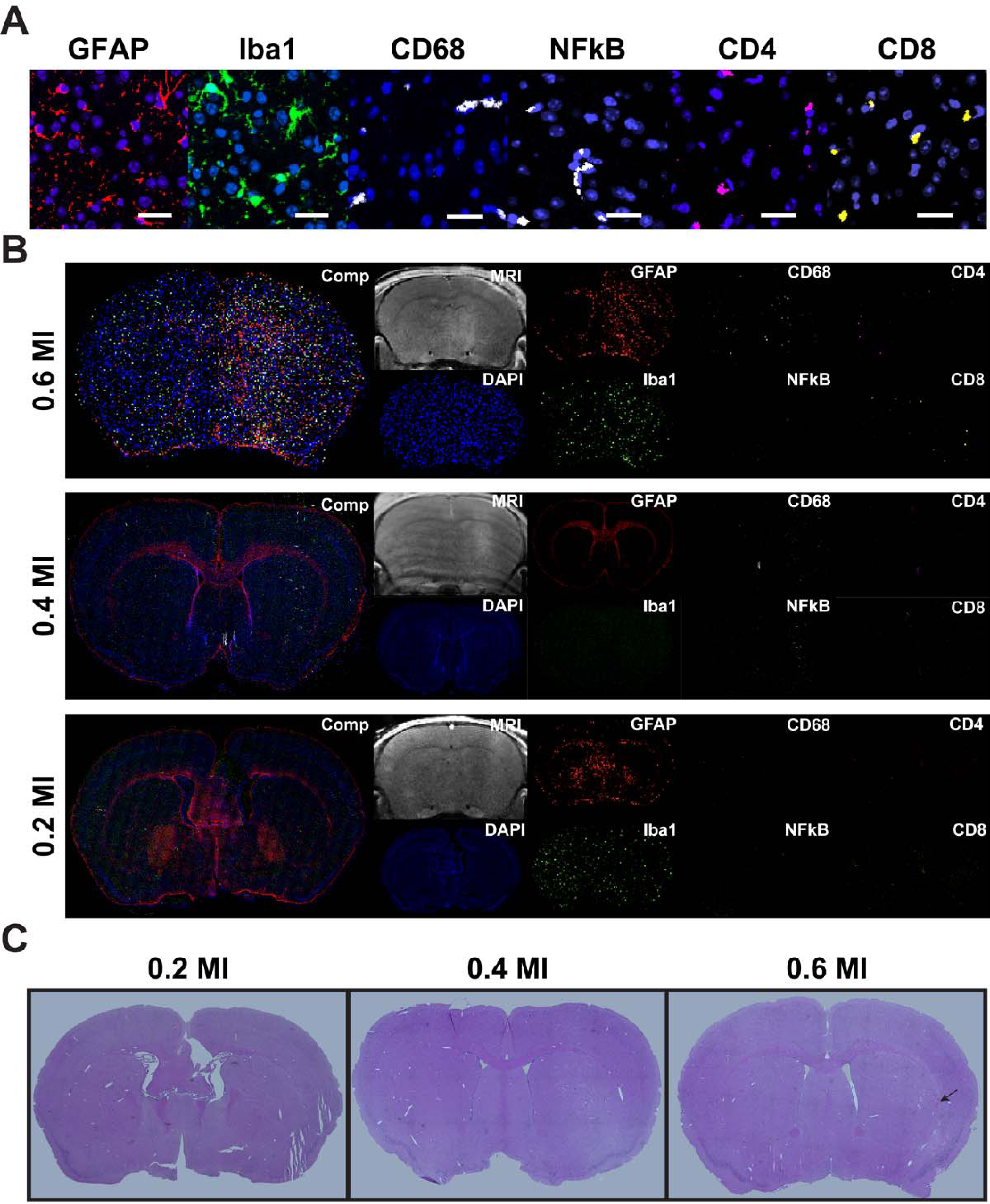
Histological Response after Blood-Brain Barrier Opening. (A) Representative images of immunohistochemistry (IHC) markers at a zoomed-in scale. Scale bar = 25 µm. (B) IHC images of three different mechanical indices (0.2, 0.4, and 0.6) at 10 MVD. A CE-T1w MRI image is provided to demonstrate the location of the blood-brain barrier (BBB) opening relative to immune cell distribution. “Comp” refers to the composite of all stains combined. (C) Hematoxylin and eosin (H&E) staining of the same doses as depicted in (B), black arrow indicates red blood cell extravasation.

### Passive Cavitation Detection Demonstrates Microbubble Activity Varies with Both MI and MVD

During each sonication, passive cavitation detection (PCD) recordings were performed to assess microbubble acoustic activity throughout the treatment. Voltage data obtained from PCD recordings were preprocessed and converted to the frequency domain (Figure 4A). As expected, the frequency content analysis revealed an increase in subharmonic and ultra-harmonic content with higher MVD and MI (Figure 4B). The presence of broadband content was observed only when the MVD exceeded 1 µL/kg at 0.6 MI (Figure 4B). The frequency content over the course of the treatments displayed a slight increase after retro-orbital injections, followed by a subsequent decline as the MBs were cleared from circulation. This phenomenon was particularly evident at the highest MVD of 40 µL/kg (Figure 4C). The average harmonic cavitation dose was calculated for all doses, and a significant linear trend was observed at all three MIs (p < 0.05, Figure 3D). Furthermore, there were no significant differences in broadband (inertial) cavitation doses between 0.2 and 0.4 MI at any MVD. However, statistically significant differences were observed at 0.6 MI with MVD higher than 1 µL/kg (Figure 4D). Transposed plots comparing harmonic and broadband cavitation dose to the MI can be found in Supplemental Figures 5A and B. These values for no FUS and no MB controls are shown in Supplemental Figure 6.

**Figure 4.**
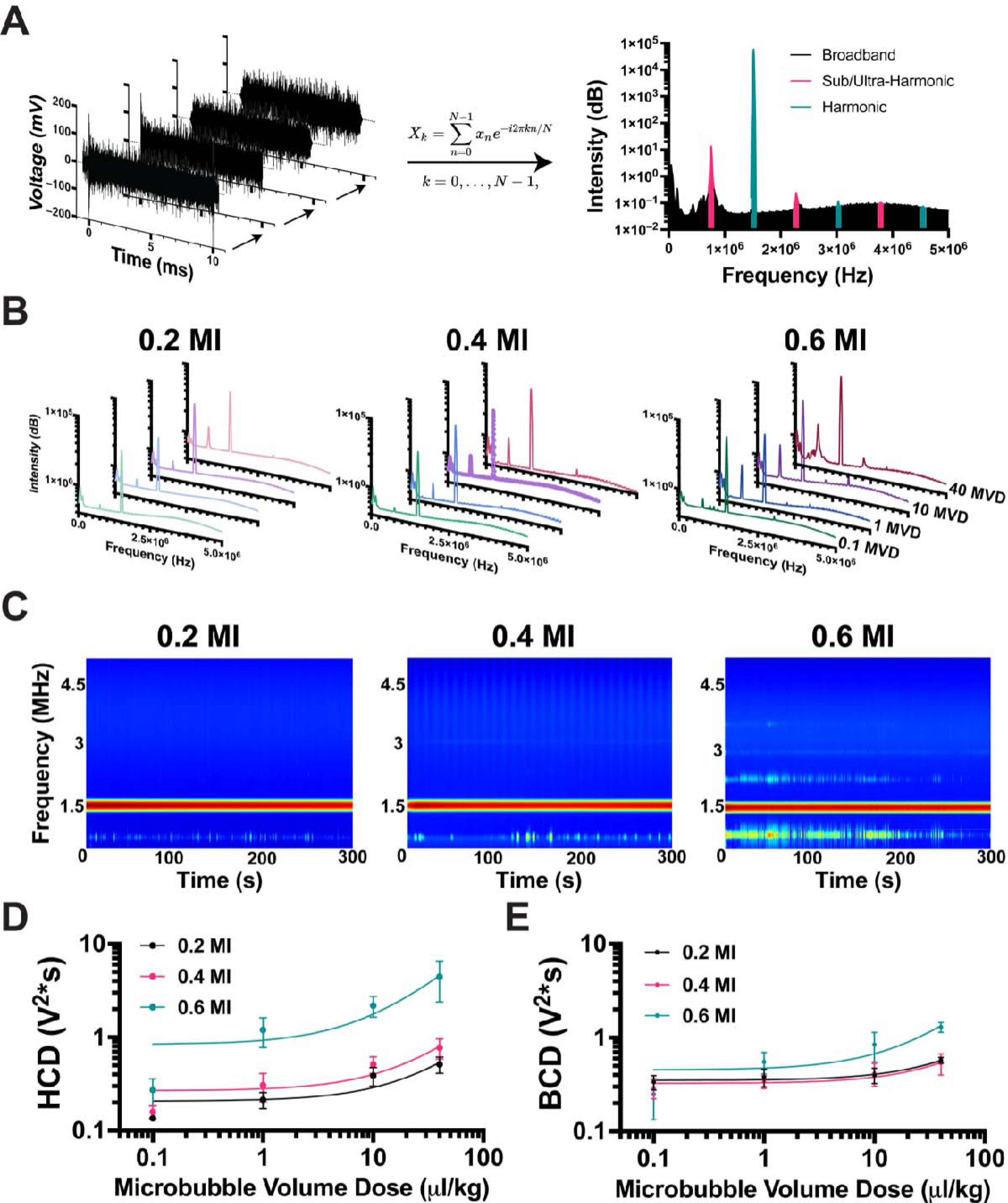
Assessment of Acoustic Response using PCD. (A) Illustration of PCD data analysis. Voltage vs time data was cropped to remove the pre-FUS signal, followed by Tukey windowing and high-frequency filtering beyond PCD sensitivity (left). The resulting signal was then converted to the frequency domain using FFT, and the area under the curve (AUC) was quantified for respective regions to determine harmonic cavitation dose (HCD) and broadband cavitation dose (BCD) (right). (B) Average FFT for each FUS pulse during a five-minute treatment at each respective MI/MVD dose. (C) Representative spectrograms for 40 MVD doses are shown throughout the sonication time (300 seconds). Quantification of harmonic cavitation dose (D) and broadband cavitation dose (E) with respect to microbubble volume dose and mechanical index (*n* = 3). Linear regression was performed for each mechanical index, resulting in R squared values for harmonic cavitation dose was 0.71, 0.74, and 0.71 for 0.2, 0.4, and 0.6 MI, respectively. The R-squared values for broadband cavitation dose were 0.73, 0.52, and 0.75 for 0.2, 0.4, and 0.6 MI, respectively. Data are presented as mean ± standard deviation.

### RNA Sequencing Indicates Differential Gene Expression Varies with MI and MVD

Six hours after MRI-guided FUS treatment, RNA was extracted from the treated brain region. Figure 5A illustrates our RNA sequencing pipeline, highlighting steps from extraction to analysis. To confirm variance between groups, all samples were initially plotted on a Principal Component Analysis (PCA) graph. Figure 5B displays the primary two components with the highest variability, accounting for 14% and 9% of the total variance. Other variability testing was conducted on our samples including UMAP and t-SNE plots found in Supplemental Figures 7A and 7B.

**Figure 5.**
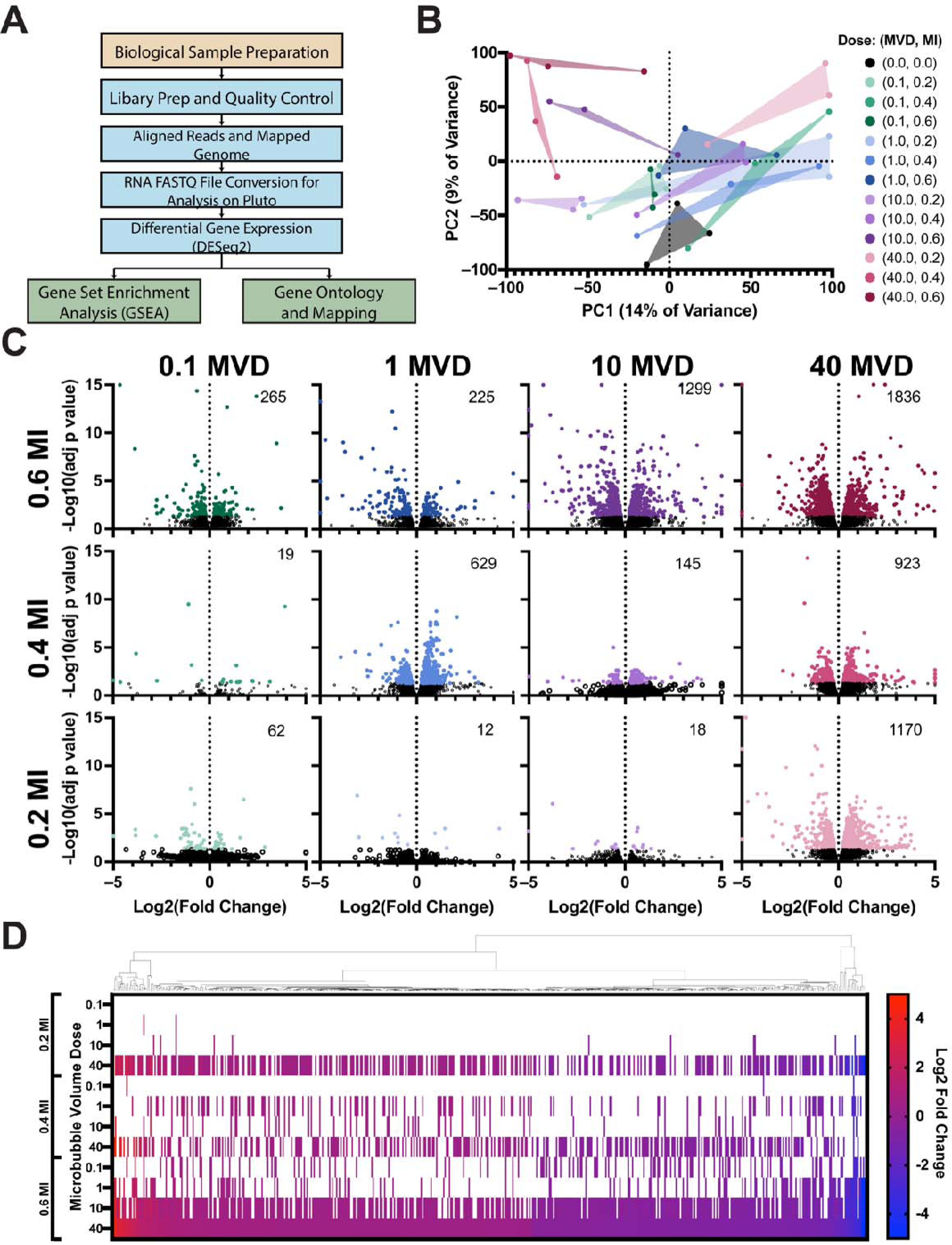
Bulk RNA Sequencing of Treated Brain Region. (A) Sequential RNA processing steps, from sample preparation to gene set enrichment analysis (GSEA) and mapping. (B) Principal component analysis (PCA) plot displaying all samples in the dataset, including different mechanical index (MI)/microbubble volume dose (MVD) combinations and isoflurane control samples. (C) Differential gene expression analysis for each MI/MVD dose. The number displayed on the top right of each plot indicates the count of significantly differentially expressed genes. All doses were compared to isoflurane control samples (*n* = 3). Among the 1,836 differentially expressed genes (DEGs) identified in the highest MI/MVD dose, 536 genes were also found in at least two other doses. (D) Heatmap illustrating the expression of the 536 genes, ranked by log2 fold change in the highest MI/MVD dose. White color represents insignificant differential expression. The Euclidean distance map is shown on top of the heatmap.

Each triplicate sample was then analyzed for differential gene expression against the no-FUS control (+Isoflurane) (Figure 5C). Overall, the differential expression of genes increased with a higher MI or MVD. The highest number of differentially expressed genes was observed in the highest MVD and MI dose (40 MVD + 0.6 MI), totaling 1836 genes; the lowest observed number was 12 at the lowest MI and MVD. To understand their significance, the 1836 differentially expressed genes from the highest dose were analyzed for similarities with other doses. Among these genes, 536 were also found in at least two other doses, indicating a specific effect of MB+FUS treatment (Figure 5D). The 536 genes were organized based on their fold change at the highest dose. As the dose was reduced in either MVD or MI, fewer genes in the set showed significant differential expression, as indicated by the white bars. Notably, the lowest dose (0.1 MVD + 0.2 MI) did not exhibit significant differential expression among the 536 genes. Furthermore, the expression levels of these 536 genes remained consistent across all doses. The genes showing higher expression levels in the highest dose (40 MVD + 0.6 MI) also demonstrated higher expression levels in all other doses. Similarly, the less expressed genes maintained their lower expression levels consistently across all doses, following a similar gradient.

### Hallmark Gene Set Enrichment Analysis Reveals Strong Inflammatory Response after BBBO

The next step in RNA analysis was to perform gene set enrichment analysis to provide a biological context for the differentially expressed genes. To cover a wide range of biological processes, we utilized the 50 hallmark gene sets from the Broad Institute of MIT and Harvard (Cambridge, Massachusetts). Figure 6A illustrates the top gene sets identified across all 12 doses. Notably, there are 12 distinct gene sets that show upregulation at the highest doses and gradually decline as the MI or MVD decreases. These twelve gene sets include, beginning with the most significant: TNFL Signaling via NFκB, Inflammatory Response, Hypoxia, Allograft Rejection, Epithelial-Mesenchymal Transition, Interferon Gamma Response, IL6 Jak Stat3 Signaling, Apoptosis, Complement, P53 Pathway, IL2 Stat5 Signaling, and Coagulation. Importantly, all these gene sets are closely associated with inflammatory responses.

**Figure 6.**
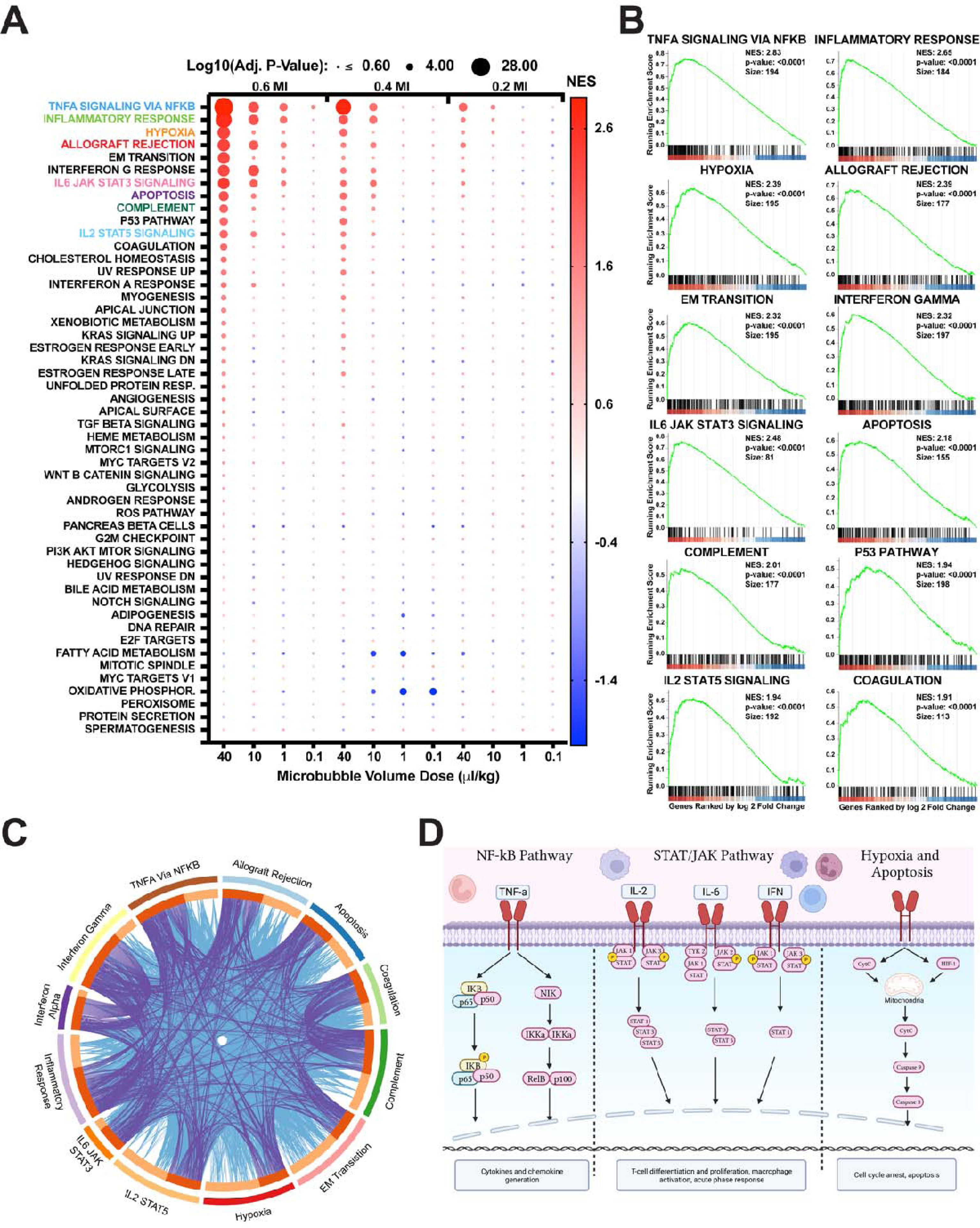
Hallmark Gene Set Enrichment Analysis. (A) Dot plot illustrating the 12 MI/MVD doses based on size (adjusted p-value) and color (normalized enrichment score). Gene sets are arranged by adjusted p-value for the highest MI/MVD dose. (B) Enrichment plot presenting the top 12 gene sets identified in (A). The x-axis displays the genes ranked by log2 fold change, with vertical ticks indicating the gene positions within the gene set. The heatmap represents gene expression, with red indicating higher expression in the first group (40 MVD 0.6 MI), and blue representing higher expression in the isoflurane control group. The green line represents the running enrichment score. (C) Overlapped Circos plot demonstrating relationships between the top 12 enriched gene sets. Purple lines indicate the presence of the same gene in different gene sets, while blue connections represent genes found in similar Gene Ontology (GO) pathways across different gene sets. (D) Schematic representation of the top enriched gene set pathways.

Enrichment plots for each of the twelve gene sets identified in the highest dose (40 MVD + 0.6 MI) are presented in Figure 6B. The high peak on the left side of the plots indicates the strong enrichment of these gene sets in our samples. It is worth noting that these gene sets exhibit close relationships with each other. To illustrate this interconnectivity, a Circos plot is shown in Figure 6C, where direct gene connections (represented by purple lines) and connections via Gene Ontology (GO) biological processes (light blue lines) are established between each gene set. Some connections show stronger relationships, such as interferon-gamma and interferon-alpha signaling, while others demonstrate weaker associations with other groups, such as Epithelial-Mesenchymal Transition.

Figure 6D provides another perspective by visualizing the major mechanisms between these gene sets through signaling molecules. Three main groups emerge from the analysis of these gene sets: the initial inflammatory response (TNFL signaling via NFκB), the major inflammatory response (which involves more chemokine and cytokine signaling), and damage-associated gene sets (related to apoptosis). Overall, the results of the gene set enrichment analysis highlight the prominent role of inflammatory responses and related pathways in the transcriptional changes observed owing to MB+FUS dose escalation.

### Blood-Brain Barrier Opening Intensity is the Best Indicator of SIR

To better understand the relationship between our parameters and BBBO or SIR, we conducted a correlation matrix analysis of all major variables identified thus far. Figure 7A presents this matrix, highlighting both strong and weak correlations. Our analysis revealed a stronger correlation between hyperintense volume and contrast enhancement in BBBO areas with MVD, as compared to the MI (0.83/0.76 and 0.41/0.48, respectively). Passive cavitation parameters exhibited relatively consistent relationships (0.46 - 0.64) with both MVD and MI. In terms of RNA expression, indicated by normalized enrichment scores (NES), correlations varied with MVD or MI. MVD showed a stronger correlation with complement, hypoxia, and TNFL signaling via NFκB (R^2^ = 0.79, 0.68, and 0.70, respectively). On the other hand, MI displayed a stronger correlation with IL2 stat5 signaling, IL6 jak stat3 signaling, interferon-gamma response, allograft rejection, and inflammatory response (R^2^ = 0.60, 0.63, 0.75, 0.61, and 0.78 respectively). Interestingly, our main finding indicated that BBBO volume and BBBO contrast enhancement were the effects most strongly associated with RNA expression, with all correlations having R^2^ values greater than 0.86 (excluding allograft rejection and interferon-gamma response). To further explore this relationship, we plotted our top-represented gene sets against BBBO volume (Figure 7B). At least half of all doses exhibited significant NES in each gene set, and each relationship demonstrated strong linear correlations.

**Figure 7.**
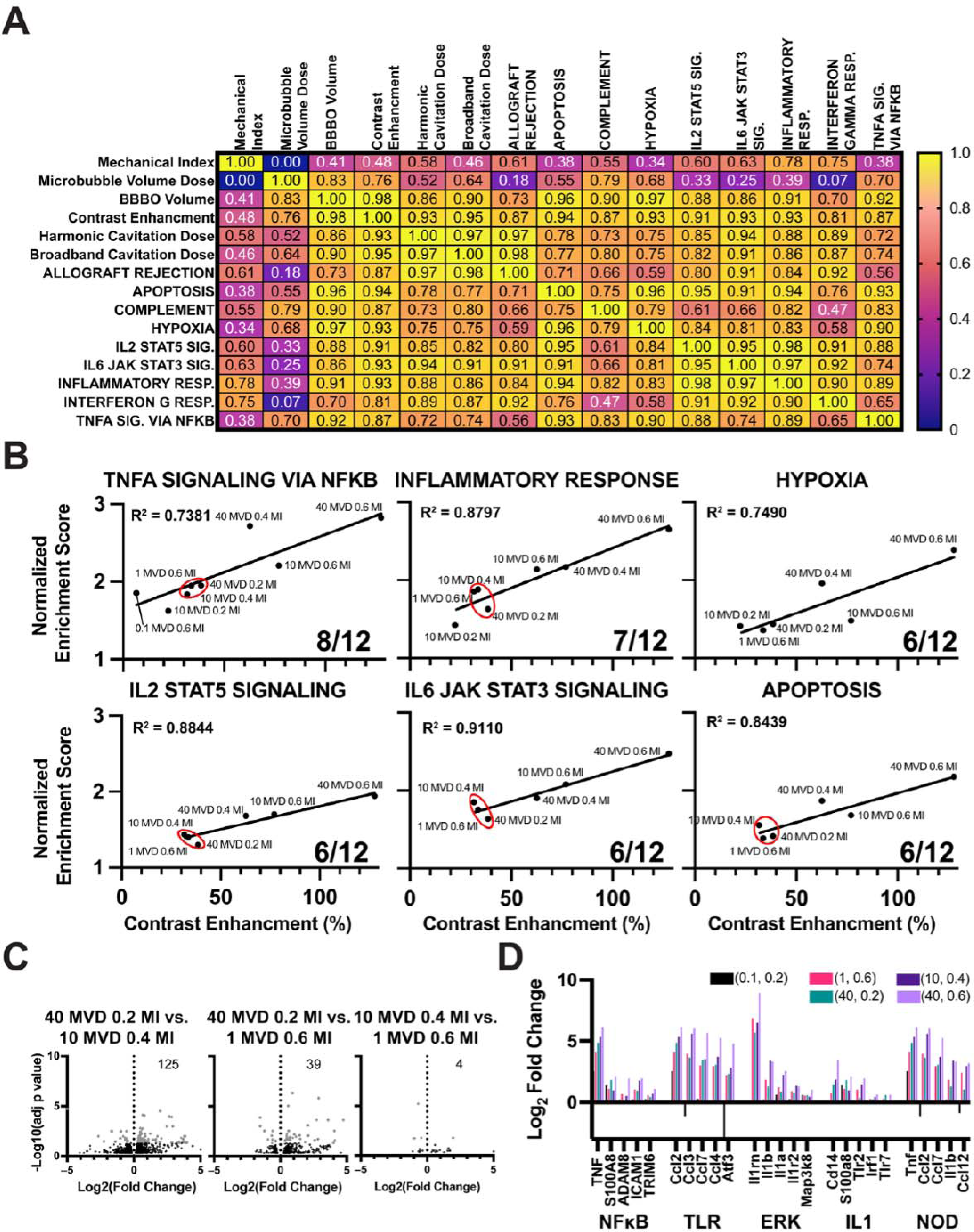
Correlation of RNA Expression and BBB Opening. (A) Correlation heatmap showing the relationship between MI and MVD doses and resulting BBBO volume, contrast enhancement (CE), cavitation doses, and RNA expression. Pearson correlation coefficient (r) values are displayed in each box. The color scale represents the magnitude of the Pearson correlation coefficient. (B) Correlation between normalized enrichment score (NES) scores and BBBO volume. Only pathways from gene set with more than six doses exhibiting significant NES scores are represented. Linear regression analysis was performed for each plot, and the corresponding R-squared value is shown in the top left corner. The plots indicate the fraction of significant doses in the bottom right. A red circle denotes a small cluster consisting of three different doses with similar BBBO volumes. (C) Volcano plot displaying the differentially expressed genes (DEGs) between doses within the small cluster (40 MVD + 0.2 MI, 10 MVD + 0.4 MI, and 1 MVD + 0.6 MI). The number of DEGs is indicated in the top right corner. (D) Bar graph representing the log2 fold change of the lowest dose (0.1 MVD + 0.2 MI), the cluster doses, and the highest dose (40 MVD + 0.6 MI). The x-axis displays five of the most influential genes within five inflammatory signaling families.

A noteworthy observation was the identification of a small cluster of three MVD/MI doses: 1 MVD + 0.6 MI, 10 MVD + 0.4 MI, and 40 MVD + 0.2 MI (highlighted by the red circle). These doses exhibited similar BBBO contrast enhancement (between 31.6 and 38.5 %) and similar NES in five out of the six gene sets. To gain further insights, we performed differential gene expression analysis between each dose (Figure 7C). The volcano plots indicated minimal differences in gene expression, with a limited number of differentially expressed genes (4-125 genes). To corroborate these findings, we examined the fold change in highly utilized genes (leading edge genes) within five inflammatory families (Figure 7D). We observed similar expression levels among the cluster doses, while their expression differed significantly from the lowest and highest doses. Overall, our results suggest that the extent of BBBO plays a pivotal role in the SIR, surpassing the influence of other parameters in isolation.

### Therapeutic Windows Can be Defined Between BBBO and SIR

As previously determined through control experiments (Supplemental Figure 1), we defined significant BBBO as a minimum of 15% contrast enhancement in a volume greater than 1 mm^3^. Figure 8A presents a dot plot depicting all doses and their corresponding BBBO volume (size) and contrast enhancement (color). Utilizing the defined thresholds, we observe a distinct region representing insignificant BBB) (blue) and significant BBBO (red).

**Figure 8.**
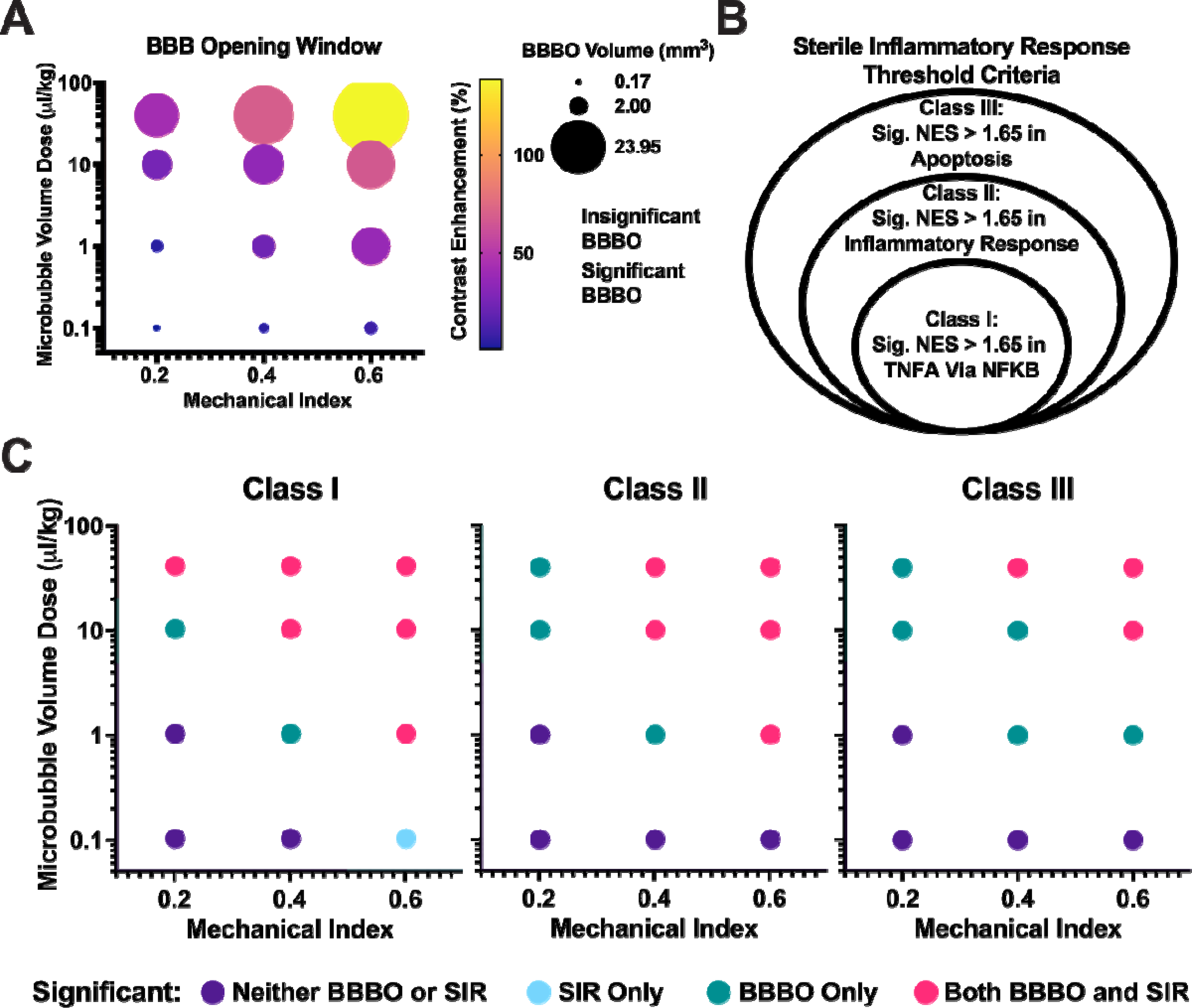
Therapeutic Window Between BBBO and Onset of Sterile Inflammatory Response (SIR). (A) Dot plot illustrating the mean blood-brain barrier opening (BBBO) in terms of volume (represented by the size of the dot) and contrast enhancement (indicated by the color of the dot). The grid is plotted based on microbubble volume dose (MVD) and mechanical index (MI). Significant BBBO is defined as 15% contrast enhancement in a 1 mm^3^ volume. The red region represents significant BBBO, while the blue region represents insignificant BBBO. (B) Classes defined for the onset of the sterile inflammatory response. Each circle represents the criteria required to meet a specific class level. (C) Dot plot displaying the regions of neither significant BBBO nor SIR (purple), BBBO only (green), SIR only (light blue), or significant BBBO and SIR (pink). Each plot represents a different class level for the onset of SIR as defined in (B).

To establish thresholds for the onset of SIR, we recognized the gradient of responses and developed different classes of sterile inflammation (Figure 8B). Each class encompasses the previous classifications, creating a hierarchical framework. Figure 8C illustrates the therapeutic window between significant BBBO and SIR. In Class I, the least strict classification, significant NES (> 1.65) in the TNFL signaling via NFκB gene set serves as the defining criterion. Within this class, a small window of significant BBB opening is observed before the onset of SIR (10 MVD + 0.2 MI and 1 MVD + 0.4 MI). Notably, this is the only instance where the onset of SIR occurs without significant BBBO (0.1 MVD + 0.6 MI).

As we progress to Class II, which is defined by the onset of significant NES (> 1.65) for both the TNFL signaling via NFκB and Inflammatory response gene sets, the therapeutic window expands. In this class, three doses exhibit significant BBBO without the onset of SIR (10-40 MVD + 0.2 MI and 1 MVD + 0.4 MI).

Finally, the most stringent classification of SIR is Class III, characterized by significant NES (> 1.65) for TNFL signaling via NFκB, Inflammatory response, and a damage-associated marker, apoptosis. Within this class, we observe five doses with significant BBBO but no onset of SIR (10-40 MVD + 0.2 MI, 1-10 MVD + 0.4 MI, and 1 MVD + 0.6 MI). Taken together, these findings delineate the therapeutic window between significant BBBO and the onset of SIR, providing valuable insights into the relationship between these two critical factors.

## Discussion

The use of focused ultrasound and microbubbles to disrupt the blood-brain barrier has gained significant attention as a promising approach for delivering therapeutics to the brain (48–52). This technique offers modularity, allowing for the manipulation of therapeutic effects. However, defining appropriate thresholds can be challenging. In our study, we focused on two key parameters: mechanical index (MI, MPa/MHz^1/2^) and microbubble volume dose (MVD, µL/kg), which have been shown to play crucial roles in determining BBB opening (18, 53). MVD is a measure of microbubble that takes into account the volume of gas that it occupies multiplied by the concentration of particles. When injected in animals, the MVD can be calculated by taking the sum of all MB volumes divided by the weight of the animals (31, 33). Prior studies performed by Song et al. showed that BBBO intensity linearly increased with increasing MVD regardless of microbubble number or size (30).

The MI combines ultrasound frequency and pressure and has been previously demonstrated to predict the threshold of BBBO (53). Consistent with previous studies (52), we observed a strong linear relationship between MI and the extent of BBBO, as measured by the signal intensity of MRI contrast enhancement. Similarly, MVD, which combines microbubble number and size, exhibited a strong linear effect on BBBO. Our findings align with previous studies (54) and confirm the significant influence of both MI and MVD on BBB opening. To explore the combined influence of MI and MVD on BBBO, we performed multiple regression analyses and observed a stronger relationship when these parameters interacted. The interaction term (MI*MVD) showed a high coefficient of determination (R^2^ = 0.9), indicating that the combined effect of MI and MVD has a greater impact on BBB opening than either parameter alone.

Passive cavitation detection provided valuable insights into microbubble behavior during the treatments. The analysis of the voltage data (34) allowed us to determine the harmonic and broadband (inertial) cavitation doses, which showed linear trends with MI and MVD. Although the R^2^ values for these relationships were slightly lower due to variability in measurements, they still demonstrated significant trends. Notably, the harmonic cavitation dose exhibited a particularly strong relationship with both BBBO volume and contrast enhancement, indicating its potential as a predictive parameter for assessing BBBO.

Determining the threshold for significant BBBO is crucial for safe and effective treatments. While various studies have investigated this threshold, the wide range of parameters and metrics used makes direct comparisons challenging. Nonetheless, many studies have reported a threshold in the range of 0.3 to 0.5 MI at different MVDs. In our study, we found the threshold for significant BBBO (>15% contrast enhancement in a volume of 1 mm^3^) to be greater than 1 MVD at 0.2 MI and greater than 0.1 MVD at 0.4 and 0.6 MI. It is noteworthy to mention that our estimates for a 15% increase in targeted signal intensity are dependent on our MRI protocol (such as the field strength of 9.4 Tesla, T1w sequence parameters, the dose and time of the injection of gadolinium chelate, etc.). The actual cortical volume of the BBBO (1 mm^3^) is the more robust parameter for the extent and limit of detection for the BBBO. When considering the combined effect of MI and MVD (MI*MVD), the threshold was found to be greater than 0.2. It is important to note that our highest dose (40 MVD + 0.6 MI) resulted in irregular tissue morphology illustrated by hyperintense and hypointense regions on T1-weighted + CE MRI. This observation suggests a potential maximum safety threshold of 40 MVD and 0.6 MI indicating that caution should be exercised at higher doses to avoid adverse tissue effects.

Differential gene expression (DEG) enables us to understand the global transcriptomic activity of a cell or tissue. However, this broad analysis excludes the specificity of which pathways are most upregulated or downregulated. Instead, the information we receive is the individual fold changes of genes. What we can extract from this type of data is that at higher doses (40 MVD + 0.6 MI) there are significantly more upregulated genes compared to the rest of the doses, meaning that more activity is occurring in that region, whether related to inflammation or not. To make sense of this data more broadly, we compared all the significant differentially expressed genes in the highest dose (40 MVD + 0.6 MI) against all the populations of DEGs in the rest of the conditions for Figure 5D. By organizing the DEG data against the 40 MVD + 0.6 MI, we can see that there is a similar trend following this order among all groups, especially when the MI and MVD decrease. Specifically, the lowest amount of differentially expressed genes is observed in 1 MVD + 0.2 MI, not the lowest dose (0.1 MVD + 0.2 MI, Fig. 5C), which is not what we expected. However, when compared against the most common DEGs, no genes were significantly up or down-regulated, indicating that the 62 DEGs are not related to inflammation or damage but rather other tangential biological processes.

To provide a biological context for the population of DEG and the pattern it exhibits we ran a gene set enrichment analysis (GSEA) with the data. The 50-hallmark gene set analysis categorized 50 of the most basic biological functions of a cell, allowing us to identify a variety of effects. All identified paths were related to inflammation and/or cellular damage, with the leading pathway being the TNFL signaling through NFκB. This supports many other studies, which identified significant upregulation of the NFκB pathway as a result of BBBO (28, 35, 55). Outside of broad inflammatory responses, one pathway that is also upregulated is the JAK/STAT and IFN pathways. These pathways have been identified in events of immune recruitment, specifically involving T-cell activation and macrophage recruitment to the site of inflammation (56, 57). Not only does this result support evidence of peripheral immune cell extravasation, but it also sheds light on the specific population of immune cells that are recruited and their specific mechanism of action. A third major event that is observed in the RNA sequencing data indicative of neuronal damage is the enrichment of genes in pathways related to hypoxia and apoptosis. Increased mechanical indices are known to be detrimental to cells, as the exerted mechanical forces exerted forces are known to cause hemorrhaging (58, 59). This has been shown in other studies (52, 53), where at higher doses, MB+FUS can cause microhemorrhages and damage that is similar to damage responses seen in traumatic brain injury (60). Additionally, altered blood flow and hypoxia have been observed as transient events accompanying BBBO, with the major mechanism implicated being capillary vessel restriction (61). Our results suggest that when microbubbles cavitate under ultrasound, they can cause a transient delay in blood perfusion, inducing local hypoxia or ischemia through hemorrhaging contributing to apoptosis. A comparison in these studies can prove that higher MVD/MI doses can mirror damage seen under these conditions. The Circos plot displays the connecting relationship between these events (Fig. 6C).

After collecting individual information about the 50-hallmark analysis, a correlation analysis was run against all the major upregulated and enriched pathways to further understand our knowledge of ultrasound and microbubble parameters as well as the most enriched pathways resulting from RNA sequencing. Interestingly, according to the results of the correlation matrix, there were characteristic qualities that were unique to MVD such as the activation of the complement system, hypoxia, and NFκB pathway via TNFL signaling. PEG, which is a reagent commonly used to improve drug longevity in the body and is found on the external shell of the microbubble, can activate the immune system through the complement system via C3b opsonization of the phosphate in the lipid headgroup (62). This phenomenon has been recognized as a potential cause of complement activation-related pseudo allergy (CARPA) and has been reported in the use of Doxil (63, 64). Surprisingly, the biggest immune pathway that has been implicated in the initiation of SIR, the NFκB pathway, was more highly correlated to MVD than MI. More in-depth work is required to define a mechanism to explain why we see this trend. On the other hand, the MI had a stronger relationship with RNA seq pathways such as Allograft rejection, IL2 STAT3, IL6 JAK STAT3, IFNγ, and inflammatory response. These five pathways are interconnected, as they represent the cascade of the inflammatory response after activation. Their stronger relationship to MI also requires a more in-depth mechanism analysis. Although these pathways correlate differently between MI and MVD, all pathways correlate most highly to BBBO volume (R^2^ > 0.70) and contrast enhancement (R^2^ > 0.81).

Normalized enrichment scores of the GSEA showed a strong linear trend when plotted against BBBO intensity (contrast enhancement %). This was evaluated among all pathways with greater than 6 significant NES. A reoccurring phenomenon of a “cluster” of MVD and MI combinations is apparent on the trendlines where 1.0 MVD + 0.6 MI and 10 MVD + 0.4 MI and 40 MVD + 0.2 MI resulted in similar intensity of opening and NES. This is seen in 5 out of the 6 graphs in Figure 7B, except Hypoxia where 10 MVD + 0.4 MI did not have a significant NES. This further supports the fact that the intensity of BBBO is the greatest indicator of bioeffects. Moreover, using a greater MVD with a lower MI can permeabilize the BBB similar to a lower MVD and higher MI, resulting in similar bioeffects. Figure 7C analyzed the differentially expressed genes in the cluster to compare the amount of differentially expressed genes and, despite having upwards of 125 differentially expressed genes down to 4, this is not considered significantly differentially expressed compared to the amount in Figure 5C with hundreds and thousands of differentially expressed genes. Figure 7D aims to further corroborate this claim by comparing the individual gene expression between 5 inflammatory pathways that are involved in sterile inflammation.

As defined previously, BBBO was considered significant when 15% contrast enhancement is achieved in a volume greater than 1 mm^3^. As seen in Figure 8A, the volume of BBBO corresponds with increased contrast enhancement. We can define a BBBO window comparing our MVDs, and MIs. Next, by comparing the RNA sequencing data from 12 experimental conditions for BBBO, differences in gene expression were analyzed and the complexity of the immune response elicited from permeabilizing the BBB was characterized. Figure 8B highlights these orders or classes by increasing the requirement of what can and should be considered a part of the sterile immune response. NFκB signaling has been defined in a plethora of studies as a hallmark of the SIR (4, 28, 35, 55). Kovacs et al. first defined this to be the major molecular mechanism behind the sterile immune response, especially after MB+FUS BBBO, with many other studies following the pursuit of further investigating the bioeffects (28, 35). Due to this, we broadly defined SIR as the activation of the NFκB pathway via TNFL signaling (NES > 1.65). Since the sterile inflammatory response is also known to prompt and recruit further immune responses, we incorporated another criterion for SIR as the addition of the significantly enriched inflammatory response (NES > 1.65). This is what we defined as Class II SIR, providing a larger window to optimize parameters. Recently, several groups have worked to operate within the SIR window after MB+FUS (65–67) without activating damage pathways (apoptosis). Our Class III window is defined as the onset of inflammatory responses without significantly activating the apoptosis pathway.

Moving forward, further investigations are warranted to expand the therapeutic windows. Firstly, exploring novel microbubble formulations or ultrasound pulsing schemes could lead to more precise and efficient BBB opening while minimizing adverse SIR-related effects. Secondly, defining the mechanisms underlying the observed bioeffects, such as inflammation, immune recruitment, and tissue damage, will provide valuable insights for the most useful class. Finally, more advanced MRI sequences can provide more insight to the microenvironment noninvasively. This includes using quantitative iron oxide-enhanced T2/ T2* weighted MRI, which could be easily incorporated into the MR guiding/ BBB kinetic protocols or taking a multi-parametric/ multi-modal imaging approach to assess inflammation, cellular damage and necrosis, as well as hypoxia using advanced non-invasive PET/MRI(68–70). All efforts will provide more tunable therapeutic windows that allow more safe and effective treatments.

In conclusion, our study demonstrates the significant influence of MVD and MI on BBBO and SIR. The combination of MI and MVD showed a stronger effect on BBBO than either parameter alone. RNA analysis revealed differential gene expression associated with inflammatory responses and immune recruitment. The study defines three therapeutic windows between significant BBBO and the onset of three classes of SIR, providing valuable guidance for safe and effective focused ultrasound-mediated drug delivery to the brain.

## Materials and Methods

### Animals

All experiments involving animals were conducted according to the regulations and policies of the Institutional Animal Care and Use Committee (IACUC) protocol 00151 in CD-1 IGS mice (strain code: 022). All mice used were female 8- to 11-wk-old and purchased from Charles River Laboratory.

### Microbubble Preparation

Lipid-coated microbubbles containing a Perfluorobutane (PFB) gas core were synthesized via sonication, as described previously by Fesitan et al. (71). Under sterile conditions, polydisperse MBs were created and collected. A single diameter (3 ± 0.5 µm) was isolated by differential centrifugation. The isolation process, including centrifugation speeds used, can be found in Fig. S8. Microbubble concentration and number- and volume-weighted size distributions were measured with a Multisizer 3 (Beckman Coulter). Microbubble concentration (c_l_, MBs/µL) versus microbubble volume (*v_l_,* µL/MB) was plotted, and the gas volume fraction (ϕ*MB*) was estimated as follows:

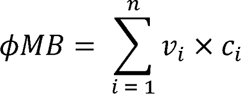

where *i* is the index of the sizing bin, 300 bins ranging from 0.7 to 18 µm in diameter. Three independent MB preparations were measured two hours before FUS treatment to confirm size distributions and concentration. Microbubble cakes were stored in the refrigerator at 4 °C for use within 1 week. Microbubbles were diluted to injection concentration within 15 minutes before injection. Supplemental Figure 9 shows MB stability 1 hour after dilution to relevant injection concentration.

### Magnetic Resonance Imaging

All animal procedures were performed under approved Institutional Animal Care and Use Committee protocols (IACUC #00151 and #0596). A Bruker BioSpec 9.4/ 20 Tesla MR Scanner (Bruker, Billerica, MA) with a mouse head RF phase-array coil was used in the Colorado Animal Imaging Shared Resource (RRID:SCR_021980). Mice were placed into in house designed MRI bed that contained stereotaxic ear bars to prevent movement of the mouse head during the transfer from MRI to focused ultrasound (FUS) system and back into the MRI scanner. Each MRI sessions consisted from a 3D-localizer a T1w MSME (Multi-Spin Multi-Echo) images were acquired in the axial plane (repetition time (TR)/echo time (TE), 720/12 ms; flip angle, 90°; number of averages, 4; field of view, 20 mm × 20 mm; matrix size, 256×256; resolution, 78 μm × 78 μm × 700 μm) was performed 12 min after intravenous injection of 0.4 mmol/kg gadobenate dimeglumine (MultiHance, Bracco, Princeton, NJ). All mice underwent high-resolution 3D T2-turboRARE (Rapid Acquisition with Relaxation Enhancement) scans (repetition time (TR)/echo time (TE), 2511/33 ms; flip angle, 90°; number of averages, 4; field of view, 20 mm × 20 mm; matrix size, 256×256; resolution, 78 μm × 78 μm × 700 μm). Mice remained stereo-tactically placed on an MRI bed and transferred to the FUS system for treatment where an intravenous injection of 0.1 mL gadobenate dimeglumine (MultiHance, Bracco, Milan, Italy) was given. All image acquisition was performed using Bruker ParaVision NEO360 v.3.3 software.

### MR Image Analysis

T1w MRI data sets were used to quantify the extent of blood-brain barrier opening using FIJI (Maryland, USA). All axial slices were analyzed by defining the contralateral hemisphere and determining the mean and standard deviation of voxel intensities. The treated hemisphere was then defined, and all voxels were found above two standard deviations of the contralateral side. The area was determined and multiplied by slice thickness (0.7 mm) to find BBBO volume. The contrast enhancement was determined by the average intensity within BBBO volume and divided by the average intensity of the control region.

### MRI-Guided FUS Treatment

The experimental setup is shown in Fig. 1D. A single element, geometrically focused transducer (frequency: 1.515 MHz, diameter: 30 mm) was driven by the RK-50 system (FUS Instruments, Toronto, Canada). A single element, geometrically focused transducer (frequency: 0.7575 MHz, diameter: 10 mm) coaxially inside the driving transducer was used for passive cavitation detection. Using the T2-weighted MR image (coronal), the center of the striatum was targeted (Fig. 1D). Ultrasound gel (Aquasonic gel, Clinton Township, MI) was placed on the mouse head confirming the lack of air bubbles. An acoustically transparent tank filled with degassed water was placed on top of the gel Fig. 1D. Microbubbles (0.1-40 μL/kg; 0.1 mL) and 0.1 mL MultiHance was injected intravenously through a retroorbital injection via 26 Ga needle. Directly after injection (within 10 seconds) FUS was applied. FUS parameters were as follows: 10 ms PL, 1 Hz PRF, 300 s treatment time, and a PNP of 0.308, 0.615, or 0.923 MPa (0.246, 0.492, or 0.738 MPa *in situ*). Voltage data from the PCD was collected during the entire FUS treatment and analyzed as previously described (34). The remaining PCD analysis was done using MATLAB (Massachusetts, USA) including the calculations of harmonic and broadband cavitation doses. Directly after FUS mice were then sent back to MRI to complete post-FUS T1-weighted imaging. Groups were divided into n = 3 for all 12 dose levels (3 MIs, 4 MVDs).

### RNA Extraction and Bulk RNA Sequencing

At 6 hours post FUS treatment mice were sacrificed via perfusion with 60 mL of ice-cold PBS. Brains were immediately dissected, and the treated site was removed and snap-frozen using liquid nitrogen. Samples were stored at -80 °C until further use. Brain samples were weighed and then immediately placed into a cell lysing buffer (Qiagen, Hilden, Germany), and homogenized for 30 seconds. RNA was isolated and purified using the RNAeasy Kit (74004, Qiagen) where all reagents were provided, and the manufacturer’s instructions were followed. Quality control and library preparation were performed through the Anschutz Genomics Core for sequencing. Poly A selected total RNA paired-end sequencing was conducted at 40 million paired reads (80 million total reads) on a NovaSEQ 6000 sequencer.

### Bulk RNA Sequencing Analysis

FASTQ files were obtained from Anschutz Genomics Core after sequencing. RNA Analysis was performed using Pluto (https://pluto.bio). Principal components analysis (PCA)(72) was calculated by applying the prcomp() R function to counts per million (CPM)-normalized values for all 40,773 targets in the experiment and samples from all groups. The data was shifted to be zero-centered. The data was scaled to have unit variance before PCA was computed. Differential expression analysis was performed comparing the groups to the isoflurane-only control group unless otherwise noted in the figure caption. Genes were filtered to include only genes with at least 3 reads counted in at least 20% of samples in any group. Differential expression analysis was then performed with the DESeq2 R package(72), which tests for differential expression based on a model using the negative binomial distribution. Log2 fold change was calculated for the comparison of the experiment to the control group. Thus, genes with a positive log2 fold change value had increased expression in experimental samples. Genes with a negative log2 fold change value had increased expression in control samples. Gene set enrichment analysis (GSEA) using the fgsea R package and the fgseaMultilevel() function (73). The log2 fold change from the experiment vs control differential expression comparison was used to rank genes. Hallmarks gene set collection from the Molecular Signatures Database (MSigDB)(74, 75) was curated using the msigdbr R package.

### Immunohistochemistry and Histology

A subset of mice (n = 7) underwent histological analysis. A representative mouse from 6 dose groups (0.2 MI at 10 MVD, 0.4MI at all MVDs, and 0.6 MI at 10 MVD, n = 1). A final mouse was only exposed to similar isoflurane levels. Six hours after treatment (if applicable) mice were sacrificed and perfused with 40 mL of 10% formalin (Thermo Fisher Scientific). Brains were immediately dissected put into a 10% formalin solution and left to shake on an orbital shaker overnight at room temperature. Primary antibody staining was done with anti-IBA1 (E4O4W, CST, Massachusetts, USA.), anti-GFAP (ab4674, Abcam, Cambridge, UK), anti-CD4 (42-0042-82, Thermo), anti-CD68 (14-0681-82, Thermo), anti-CD8 (MA514548, Thermo), anti-CD44 (701406, Thermo), anti-NFκB-p65 (51-0500, Thermo. Samples were incubated overnight at 4 °C. Signals were detected using three fluorescently labeled secondary antibodies which include: G_anti_AF594 (rat, A-11007, Thermo), G_anti_ AF488 (chicken, A-11042, Thermo) and G_anti_AF488 (rabbit, A-11008, Thermo). Expected cell types for each stain can be found in supplemental table 2. Microscope slides were imaged on a spinning disk confocal microscope (Nikon, Tokyo, Japan) at 20x. Quantification of images was done in FIJI (NIH, Maryland, USA) where an Analyze Particles function was used to determine the location of nuclei (DAPI staining) and compared the location of fluorescent signal from antibodies. If the locations matched at more than 5 pixels, the cell was counted, and the location was noted. Each slice was analyzed at all locations within. Each sample was also stained with Hematoxylin/Eosin and Luxol Fast Blue to determine the effect on tissue morphology. These two stains were imaged on a Nikon brightfield microscope.

### Statistical Analysis

All data collected is presented as mean ± SD. No preprocessing was done to the data except for voltage data collected from the PCD. PCD data were preprocessed as described in Martinez et al(34). All statistical analysis was completed in Prism 9 (GraphPad, California, USA). Star representations of p-values are indicated in captions and less than 0.05 was indicative of statistical significance. An unpaired Student’s t-test and ANOVA/multiple comparisons were used to compare two groups and larger comparisons respectively. The false discovery rate (FDR) method was applied for multiple testing correction (76). An adjusted p-value of 0.01 was used as the threshold for statistical significance.

## Acknowledgments

This work is supported by the University of Colorado Cancer Center Support Grant (P30CA046934, NJS) and NIH S10 Shared Resource grant (S10OD023485, NJS). We would like to thank the histology core at the Anschutz medical campus, in particular Laura Hoagin, for conducting all slicing and staining of our IHC samples. The Genomics core, in particular Schuyler Lee, was extremely helpful with not only the sequencing of the RNA but also their guidance throughout the bioinformatics process. Parts of Figures 1 and 5 were created using Biorender.com.

## References

1. K. Hynynen, N. McDannold, N. Vykhodtseva, F. A. Jolesz, Noninvasive MR imaging-guided focal opening of the blood-brain barrier in rabbits. Radiology 220, 640–646 (2001).

2. S.-K. Wu, C.-L. Tsai, Y. Huang, K. Hynynen, Focused Ultrasound and Microbubbles-Mediated Drug Delivery to Brain Tumor. Pharmaceutics 13, 15 (2020).

3. K. Mitusova, et al., Overcoming the blood–brain barrier for the therapy of malignant brain tumor: current status and prospects of drug delivery approaches. J. Nanobiotechnology 20, 412 (2022).

4. O. Jung, et al., Neuroinflammation associated with ultrasound-mediated permeabilization of the blood–brain barrier. Trends Neurosci. 45, 459–470 (2022).

5. N. Todd, et al., Secondary effects on brain physiology caused by focused ultrasound-mediated disruption of the blood–brain barrier. J. Controlled Release 324, 450–459 (2020).

6. C. Menaceur, F. Gosselet, L. Fenart, J. Saint-Pol, The Blood–Brain Barrier, an Evolving Concept Based on Technological Advances and Cell–Cell Communications. Cells 11, 133 (2021).

7. B. W. Chow, C. Gu, The Molecular Constituents of the Blood–Brain Barrier. Trends Neurosci. 38, 598–608 (2015).

8. W. M. Pardridge, The blood-brain barrier: Bottleneck in brain drug development. NeuroRX 2, 3–14 (2005).

9. A.-C. Luissint, C. Artus, F. Glacial, K. Ganeshamoorthy, P.-O. Couraud, Tight junctions at the blood brain barrier: physiological architecture and disease-associated dysregulation. Fluids Barriers CNS 9, 23 (2012).

10. J. A. Siegenthaler, F. Sohet, R. Daneman, ‘Sealing off the CNS’: cellular and molecular regulation of blood–brain barriergenesis. Curr. Opin. Neurobiol. 23, 1057–1064 (2013).

11. W. A. Mills, et al., Astrocyte plasticity in mice ensures continued endfoot coverage of cerebral blood vessels following injury and declines with age. Nat. Commun. 13, 1794 (2022).

12. L. S. Brown, et al., Pericytes and Neurovascular Function in the Healthy and Diseased Brain. Front. Cell. Neurosci. 13, 282 (2019).

13. H. Kubotera, et al., Astrocytic endfeet re-cover blood vessels after removal by laser ablation. Sci. Rep. 9, 1263 (2019).

14. A. L. Klibanov, Targeted delivery of gas-filled microspheres, contrast agents for ultrasound imaging. Adv. Drug Deliv. Rev. 37, 139–157 (1999).

15. E. C. Unger, E. Hersh, M. Vannan, T. McCreery, Gene delivery using ultrasound contrast agents. Echocardiogr. Mt. Kisco N 18, 355–361 (2001).

16. K. Ferrara, R. Pollard, M. Borden, Ultrasound Microbubble Contrast Agents: Fundamentals and Application to Gene and Drug Delivery. Annu. Rev. Biomed. Eng. 9, 415–447 (2007).

17. M. A. Borden, K.-H. Song, Reverse engineering the ultrasound contrast agent. Adv. Colloid Interface Sci. 262, 39–49 (2018).

18. R. E. Apfel, C. K. Holland, Gauging the likelihood of cavitation from short-pulse, low-duty cycle diagnostic ultrasound. Ultrasound Med. Biol. 17, 179–185 (1991).

19. D. L. Miller, Overview of experimental studies of biological effects of medical ultrasound caused by gas body activation and inertial cavitation. Prog. Biophys. Mol. Biol. 93, 314–330 (2007).

20. J. J. Choi, M. Pernot, S. A. Small, E. E. Konofagou, Noninvasive, transcranial and localized opening of the blood-brain barrier using focused ultrasound in mice. Ultrasound Med. Biol. 33, 95–104 (2007).

21. M. A. O’Reilly, K. Hynynen, Blood-brain barrier: real-time feedback-controlled focused ultrasound disruption by using an acoustic emissions-based controller. Radiology 263, 96–106 (2012).

22. M. Bismuth, et al., Acoustically Detonated Microbubbles Coupled with Low Frequency Insonation: Multiparameter Evaluation of Low Energy Mechanical Ablation. Bioconjug. Chem. 33, 1069–1079 (2022).

23. Y. Yang, et al., Promoting the effect of microbubble-enhanced ultrasound on hyperthermia in rabbit liver. *J*. Med. Ultrason. 49, 133–142 (2022).

24. Y.-S. Tung, F. Vlachos, J. A. Feshitan, M. A. Borden, E. E. Konofagou, The mechanism of interaction between focused ultrasound and microbubbles in blood-brain barrier opening in mice. J. Acoust. Soc. Am. 130, 3059–3067 (2011).

25. D. McMahon, M. A. O’Reilly, K. Hynynen, Therapeutic Agent Delivery Across the Blood– Brain Barrier Using Focused Ultrasound. Annu. Rev. Biomed. Eng. 23, 89–113 (2021).

26. C. Poon, D. McMahon, K. Hynynen, Noninvasive and targeted delivery of therapeutics to the brain using focused ultrasound. Neuropharmacology 120, 20–37 (2017).

27. E. E. Konofagou, Optimization of the Ultrasound-Induced Blood-Brain Barrier Opening. Theranostics 2, 1223–1237 (2012).

28. D. McMahon, K. Hynynen, Acute Inflammatory Response Following Increased Blood-Brain Barrier Permeability Induced by Focused Ultrasound is Dependent on Microbubble Dose. Theranostics 7, 3989–4000 (2017).

29. K. Gandhi, A. Barzegar-Fallah, A. Banstola, S. B. Rizwan, J. N. J. Reynolds, Ultrasound-Mediated Blood–Brain Barrier Disruption for Drug Delivery: A Systematic Review of Protocols, Efficacy, and Safety Outcomes from Preclinical and Clinical Studies. Pharmaceutics 14, 833 (2022).

30. K.-H. Song, et al., Microbubble gas volume: A unifying dose parameter in blood-brain barrier opening by focused ultrasound. Theranostics 7, 144–152 (2017).

31. K.-H. Song, B. K. Harvey, M. A. Borden, State-of-the-art of microbubble-assisted blood-brain barrier disruption. Theranostics 8, 4393–4408 (2018).

32. S. Sirsi, J. Feshitan, J. Kwan, S. Homma, M. Borden, Effect of microbubble size on fundamental mode high frequency ultrasound imaging in mice. Ultrasound Med. Biol. 36, 935– 948 (2010).

33. J. A. Navarro-Becerra, K.-H. Song, P. Martinez, M. A. Borden, Microbubble Size and Dose Effects on Pharmacokinetics. ACS Biomater. Sci. Eng. 8, 1686–1695 (2022).

34. P. Martinez, N. Bottenus, M. Borden, Cavitation Characterization of Size-Isolated Microbubbles in a Vessel Phantom Using Focused Ultrasound. Pharmaceutics 14, 1925 (2022).

35. Z. I. Kovacs, et al., Disrupting the blood–brain barrier by focused ultrasound induces sterile inflammation. Proc. Natl. Acad. Sci. 114 (2017).

36. T. Gong, L. Liu, W. Jiang, R. Zhou, DAMP-sensing receptors in sterile inflammation and inflammatory diseases. Nat. Rev. Immunol. 20, 95–112 (2020).

37. K. L. Rock, E. Latz, F. Ontiveros, H. Kono, The Sterile Inflammatory Response. Annu. Rev. Immunol. 28, 321–342 (2010).

38. C.-J. Chen, et al., Identification of a key pathway required for the sterile inflammatory response triggered by dying cells. Nat. Med. 13, 851–856 (2007).

39. G. Y. Chen, G. Nuñez, Sterile inflammation: sensing and reacting to damage. Nat. Rev. Immunol. 10, 826–837 (2010).

40. N. Feldman, A. Rotter-Maskowitz, E. Okun, DAMPs as mediators of sterile inflammation in aging-related pathologies. Ageing Res. Rev. 24, 29–39 (2015).

41. P. A. Keyel, How is inflammation initiated? Individual influences of IL-1, IL-18 and HMGB1. Cytokine 69, 136–145 (2014).

42. Y. Chen, M. N. Yousaf, W. Z. Mehal, Role of sterile inflammation in fatty liver diseases. Liver Res. 2, 21–29 (2018).

43. M. Z. Ratajczak, et al., Sterile Inflammation of Brain, due to Activation of Innate Immunity, as a Culprit in Psychiatric Disorders. Front. Psychiatry 9, 60 (2018).

44. K. Otani, T. Shichita, Cerebral sterile inflammation in neurodegenerative diseases. Inflamm. Regen. 40, 28 (2020).

45. S. Sinharay, et al., In vivo imaging of sterile microglial activation in rat brain after disrupting the blood-brain barrier with pulsed focused ultrasound: [18F]DPA-714 PET study. J. Neuroinflammation 16, 155 (2019).

46. C. Poon, C. Pellow, K. Hynynen, Neutrophil recruitment and leukocyte response following focused ultrasound and microbubble mediated blood-brain barrier treatments. Theranostics 11, 1655–1671 (2021).

47. L. Marchetti, B. Engelhardt, Immune cell trafficking across the blood-brain barrier in the absence and presence of neuroinflammation. *Vasc*. Biol. 2, H1–H18 (2020).

48. S. Alli, et al., Brainstem blood brain barrier disruption using focused ultrasound: A demonstration of feasibility and enhanced doxorubicin delivery. J. Controlled Release 281, 29–41 (2018).

49. C. D. Arvanitis, et al., Mechanisms of enhanced drug delivery in brain metastases with focused ultrasound-induced blood–tumor barrier disruption. Proc. Natl. Acad. Sci. 115 (2018).

50. C. Bing, et al., Characterization of different bubble formulations for blood-brain barrier opening using a focused ultrasound system with acoustic feedback control. Sci. Rep. 8, 7986 (2018).

51. C. Chaves, et al., Characterization of the Blood–Brain Barrier Integrity and the Brain Transport of SN-38 in an Orthotopic Xenograft Rat Model of Diffuse Intrinsic Pontine Glioma. Pharmaceutics 12, 399 (2020).

52. P.-C. Chu, et al., Focused ultrasound-induced blood-brain barrier opening: association with mechanical index and cavitation index analyzed by dynamic contrast-enhanced magnetic-resonance imaging. Sci. Rep. 6, 1–13 (2016).

53. N. McDannold, N. Vykhodtseva, K. Hynynen, Blood-Brain Barrier Disruption Induced by Focused Ultrasound and Circulating Preformed Microbubbles Appears to Be Characterized by the Mechanical Index. Ultrasound Med. Biol. 34, 834–840 (2008).

54. K.-H. Song, et al., Microbubble gas volume: A unifying dose parameter in blood-brain barrier opening by focused ultrasound. Theranostics 7, 144–152 (2017).

55. H. J. Choi, et al., The new insight into the inflammatory response following focused ultrasound-mediated blood–brain barrier disruption. Fluids Barriers CNS 19, 103 (2022).

56. X. Hu, J. Li, M. Fu, X. Zhao, W. Wang, The JAK/STAT signaling pathway: from bench to clinic. Signal Transduct. Target. Ther. 6, 402 (2021).

57. M. Jain, et al., Role of JAK/STAT in the Neuroinflammation and its Association with Neurological Disorders. Ann. Neurosci. 28, 191–200 (2021).

58. C.-H. Fan, et al., Detection of Intracerebral Hemorrhage and Transient Blood-Supply Shortage in Focused-Ultrasound-Induced Blood–Brain Barrier Disruption by Ultrasound Imaging. Ultrasound Med. Biol. 38, 1372–1382 (2012).

59. H.-C. Tsai, et al., Safety evaluation of frequent application of microbubble-enhanced focused ultrasound blood-brain-barrier opening. Sci. Rep. 8, 17720 (2018).

60. Z. I. Kovacs, et al., MRI and histological evaluation of pulsed focused ultrasound and microbubbles treatment effects in the brain. Theranostics 8, 4837–4855 (2018).

61. P.-C. Chu, et al., Neuromodulation accompanying focused ultrasound-induced blood-brain barrier opening. Sci. Rep. 5, 15477 (2015).

62. N. d’Avanzo, et al., Immunogenicity of Polyethylene Glycol Based Nanomedicines: Mechanisms, Clinical Implications and Systematic Approach. Adv. Ther. 3, 1900170 (2020).

63. C. C. Chen, M. A. Borden, The role of poly(ethylene glycol) brush architecture in complement activation on targeted microbubble surfaces. Biomaterials 32, 6579–6587 (2011).

64. J. Szebeni, Complement activation-related pseudoallergy: A new class of drug-induced acute immune toxicity. Toxicology 216, 106–121 (2005).

65. H.-L. Liu, et al., Low-pressure pulsed focused ultrasound with microbubbles promotes an anticancer immunological response. J. Transl. Med. 10, 221 (2012).

66. C. T. Curley, N. D. Sheybani, T. N. Bullock, R. J. Price, Focused Ultrasound Immunotherapy for Central Nervous System Pathologies: Challenges and Opportunities. Theranostics 7, 3608–3623 (2017).

67. P. Bathini, et al., Acute Effects of Focused Ultrasound-Induced Blood-Brain Barrier Opening on Anti-Pyroglu3 Abeta Antibody Delivery and Immune Responses. Biomolecules 12, 951 (2022).

68. N. J. Serkova, Nanoparticle-Based Magnetic Resonance Imaging on Tumor-Associated Macrophages and Inflammation. Front. Immunol. 8, 590 (2017).

69. N. J. Serkova, et al., Renal Inflammation: Targeted Iron Oxide Nanoparticles for Molecular MR Imaging in Mice. Radiology 255, 517–526 (2010).

70. G. Wang, N. J. Serkova, E. V. Groman, R. I. Scheinman, D. Simberg, Feraheme (Ferumoxytol) Is Recognized by Proinflammatory and Anti-inflammatory Macrophages via Scavenger Receptor Type AI/II. Mol. Pharm. 16, 4274–4281 (2019).

71. J. A. Feshitan, C. C. Chen, J. J. Kwan, M. A. Borden, Microbubble size isolation by differential centrifugation. J. Colloid Interface Sci. 329, 316–324 (2009).

72. M. I. Love, W. Huber, S. Anders, Moderated estimation of fold change and dispersion for RNA-seq data with DESeq2. Genome Biol. 15, 550 (2014).

73. G. Korotkevich, et al., “Fast gene set enrichment analysis” (Bioinformatics, 2016) 10.1101/060012 (June 24, 2023).

74. A. Liberzon, et al., The Molecular Signatures Database Hallmark Gene Set Collection. Cell Syst. 1, 417–425 (2015).

75. A. Subramanian, et al., Gene set enrichment analysis: A knowledge-based approach for interpreting genome-wide expression profiles. Proc. Natl. Acad. Sci. 102, 15545–15550 (2005).

76. Y. Benjamini, Y. Hochberg, Controlling the False Discovery Rate: A Practical and Powerful Approach to Multiple Testing. J. R. Stat. Soc. Ser. B Methodol. 57, 289–300.

