## Supplemental Materials for "Comprehensive Assessment of Blood-Brain Barrier Opening and Sterile Inflammatory Response: Unraveling the Therapeutic Window"

**The PDF file includes:**

Table 1: **Microbubble formulation characterization.**
Fig. 1: **Quantification of Pre T1w MR Images.**
Fig. 2: **Transpose of BBBO Volume and Contrast Enhancement.**
Fig. 3: **FIJI Circle and Round Scores.**
Fig. 4: **IHC Images (Luxol Fast Blue and CD44).**
Fig. 5: **Transpose of Harmonic and Broadband Cavitation Doses.**
Fig. 6: **Passive Cavitation Detection Controls.**
Fig. 7: **UMAP & t-SNE of RNA Sequencing Groups.**
Fig. 8: **Size Isolation Process of Microbubbles.**
Fig. 9: **Microbubble Stability.**
Table 2: **IHC Stains Used and Cell Types Expected.**

References

**Supplementary Table 1. Microbubble formulation characterization**. The table shows the mean diameter, d10 - d90 (where 10 - 90 percent of all MBs fall below this diameter), in the number-weighted graph. The mean diameter for the volume-weighted distribution is also shown. The average gas volume fraction ($\phi MB$ ) for the average concentration is given in the rightmost column. All values were averaged over 24 separate measurements (8 samples).

| **Size Distribution Characterization** | | | | | | | | | |
|---|---|---|---|---|---|---|---|---|---|
| **Number % Diameter Parameters (μm)** | | | | | | **Volume % Mean Diameter (μm)** | **Mean Volume (μm$^3$)** | **Mean Conc. (MBs/mL)** | ΦMB (μL/mL) |
| **Mean** | **$d_{10}$** | **$d_{25}$** | **$d_{50}$** | **$d_{75}$** | **$d_{90}$** | | | | |
| 3.32 ± 0.15 | 2.53 ± 0.07 | 2.90 ± 0.11 | 3.30 ± 0.17 | 3.74 ± 0.20 | 4.16 ± 0.22 | 3.73 ± 0.33 | 17 ± 2.5 | 1.1E10 ± 2.4E9 | 187 ± 27.5 |

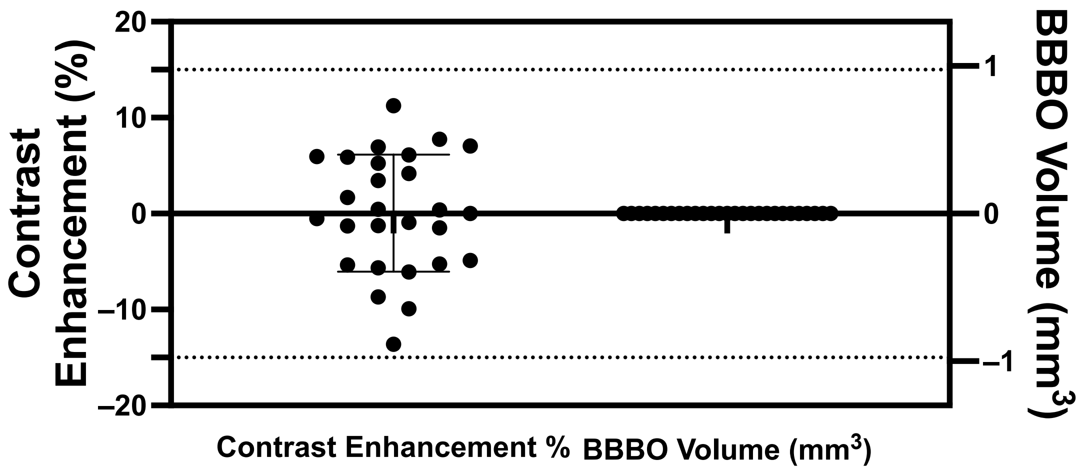


**Supplementary Fig. 1: Quantification of Pre T1w MR Images.** Pre T1w MR images were used to confirm intact BBB prior to FUS/MB treatments. Contrast enhancement was determined between each hemisphere. Dotted lines represent used thresholds for significant BBB opening (15% and 1 $mm^3$). Data is presented as mean ± standard deviation (*n* = 27).

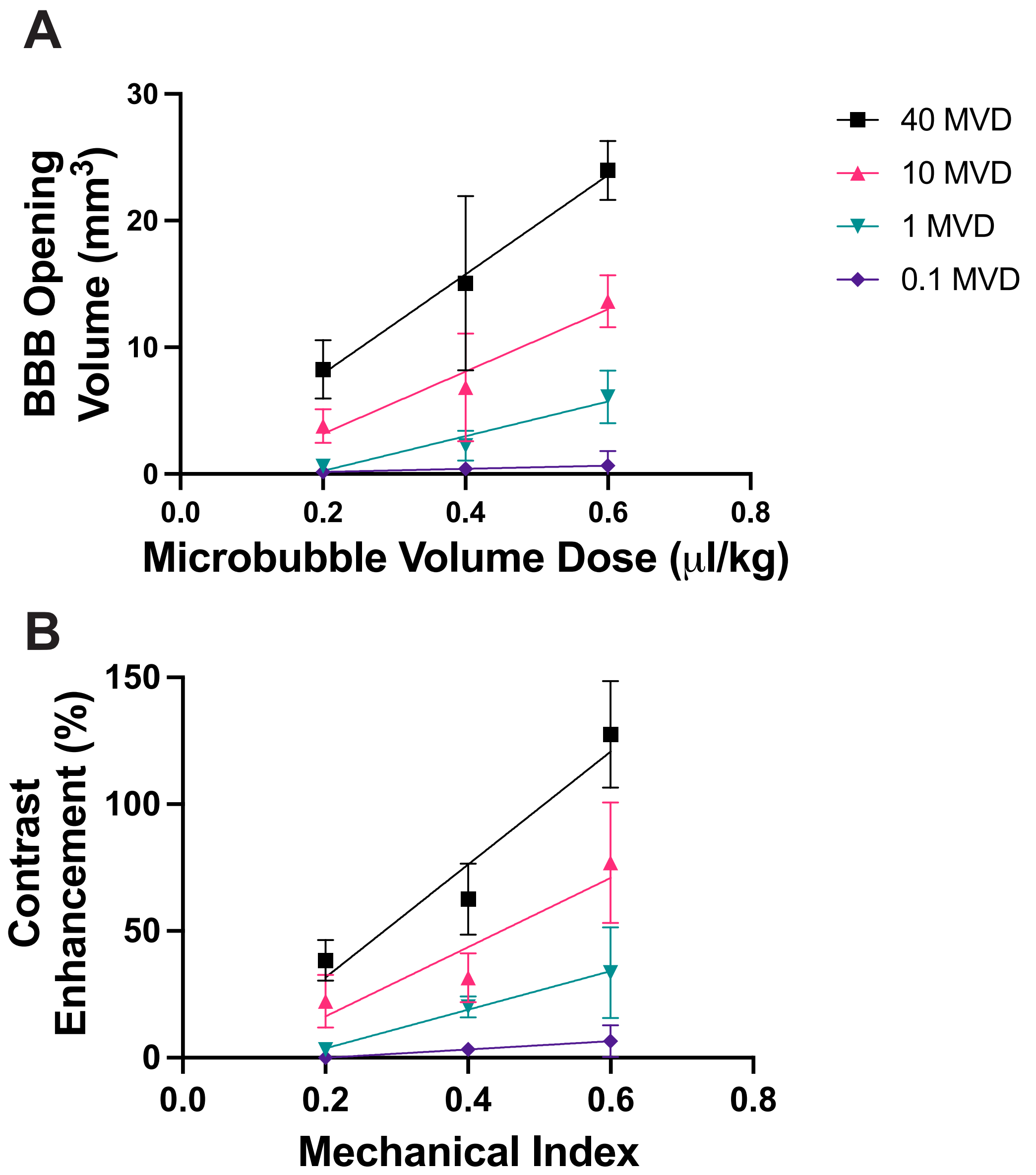


**Supplementary Fig. 2: Transpose of BBBO Volume and Contrast Enhancement.** (A) Quantification of BBBO volume (*n* = 3). Linear regression was performed for each mechanical index, resulting in R squared values of 0.11, 0.76, 0.72, and 0.76 for 0.1, 1, 10, and 40 MVD, respectively. (B) Quantification of BBBO contrast enhancement (*n* = 3). Linear regression was conducted for each mechanical index, resulting in R squared values of 0.40, 0.67, 0.67, and 0.84 for 0.1, 1, 10, and 40 MVD, respectively. Data is presented as mean ± standard deviation.

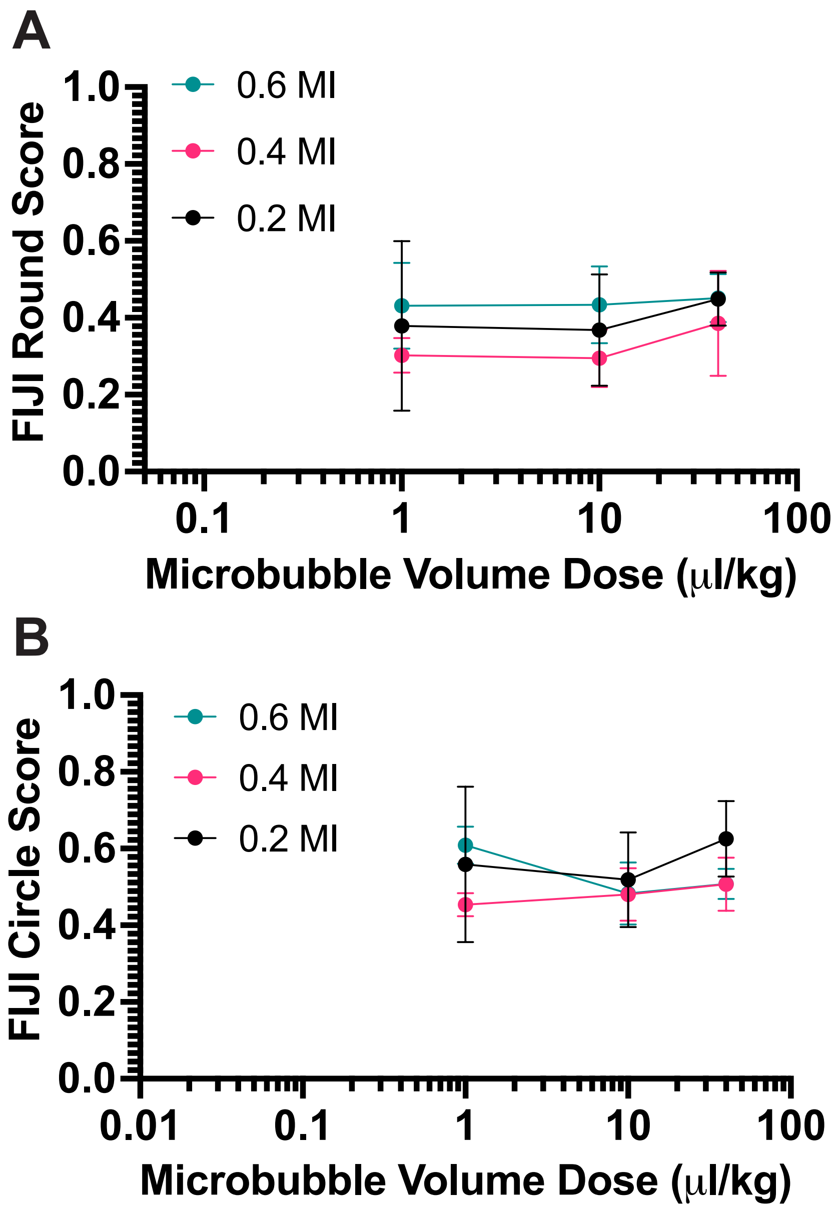


**Supplementary Fig. 3: FIJI Circle and Round Scores.** Quantification of BBBO volume's FIJI round (A) and circle (B) score (*n* = 3). There is no significant difference between any groups. Data is presented as mean ± standard deviation.

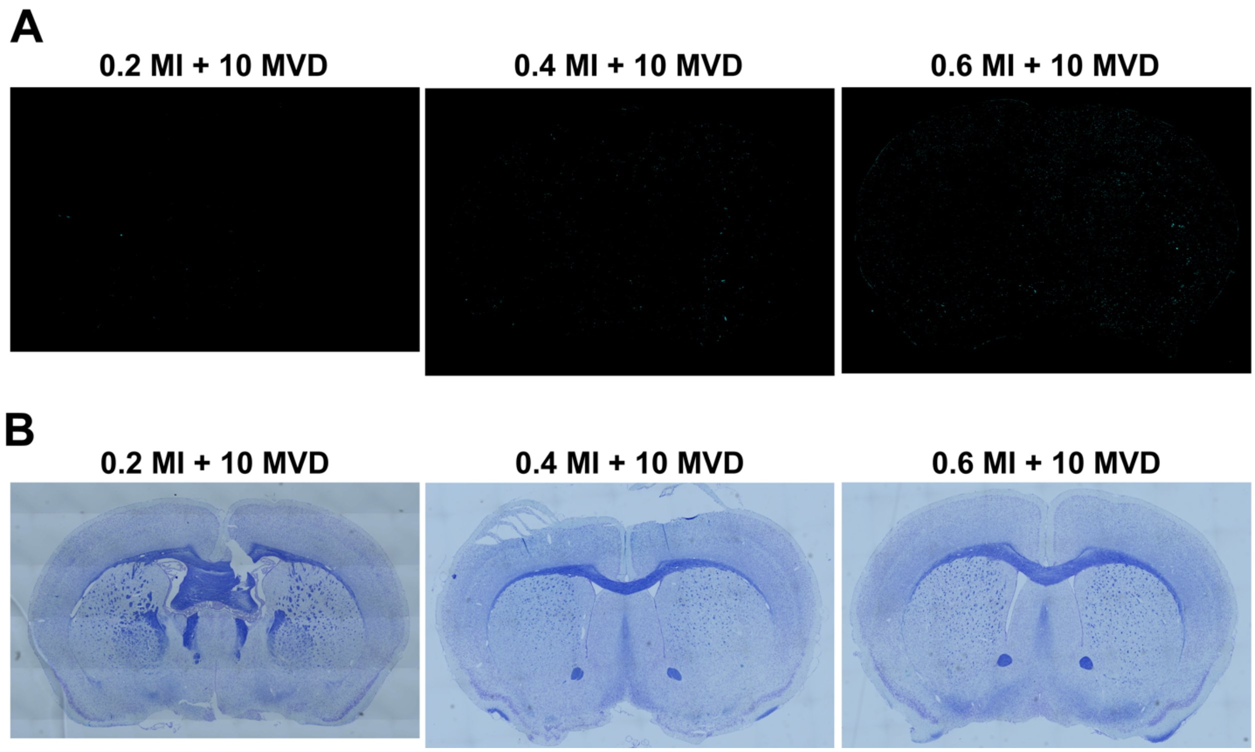


**Supplementary Fig. 4: IHC Images (Luxol Fast Blue and CD44).** IHC images of CD44 (A) and LFB (B) at three different mechanical indices (0.2, 0.4, and 0.6) at 10 MVD.

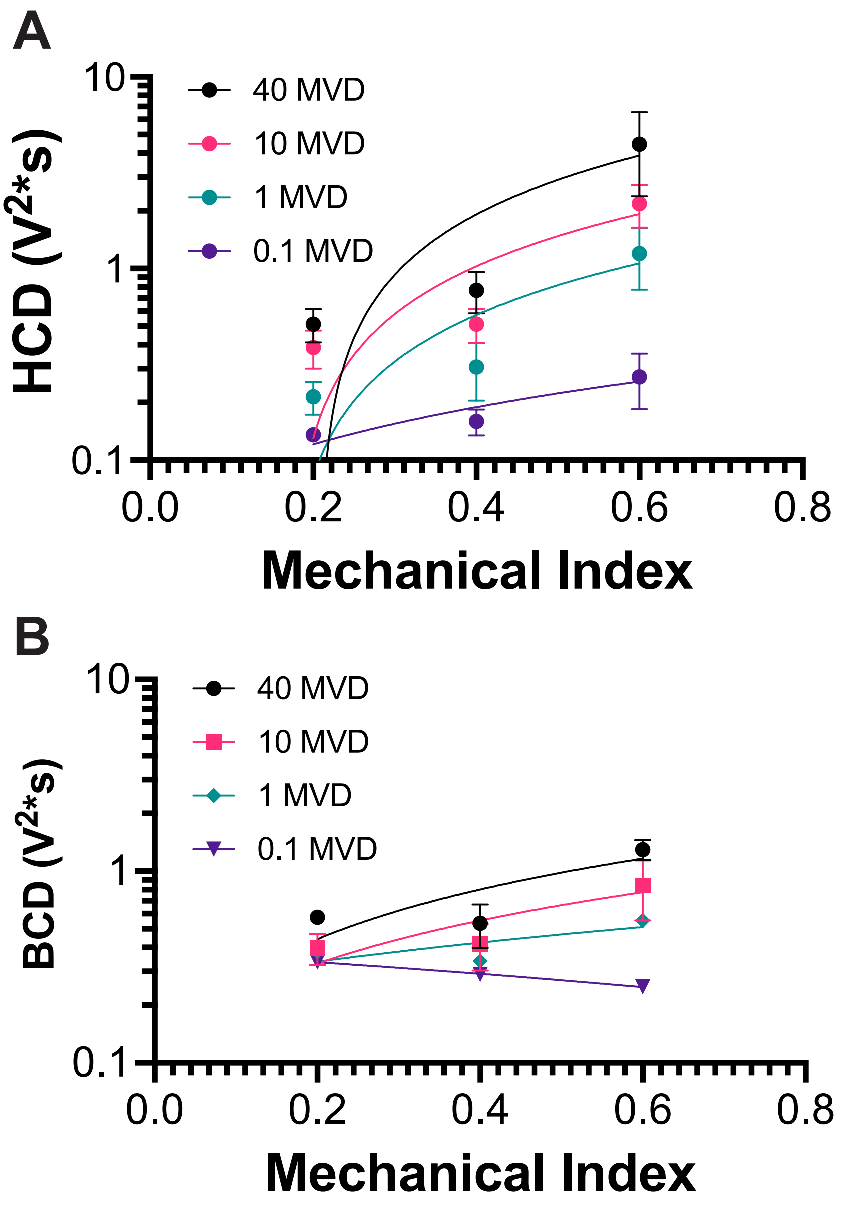


**Supplementary Fig. 5: Transpose of Harmonic and Broadband Cavitation Doses.** Harmonic (A) and broadband cavitation dose (B) with respect to microbubble volume dose and mechanical index (*n* = 3). Linear regression was performed for each mechanical index, resulting in R squared values for harmonic cavitation dose was 0.58, 0.67, 0.73, and 0.62 for 0.1, 1, 10, and 40 MVD, respectively. The R-squared values for broadband cavitation dose were 0.21, 0.33, 0.51, and 0.66 for 0.1, 1, 10, and 40 MVD, respectively. Data is presented as mean ± standard deviation.

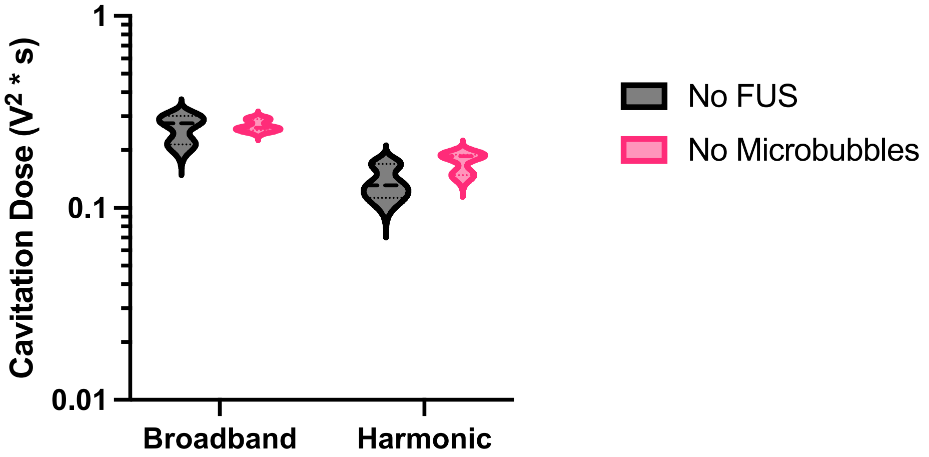


**Supplementary Fig. 6: Passive Cavitation Detection Controls.** The passive cavitation detection without FUS application or microbubbles. Both harmonic (right) and broadband (left) cavitation doses are shown (*n* = 3).

**A**

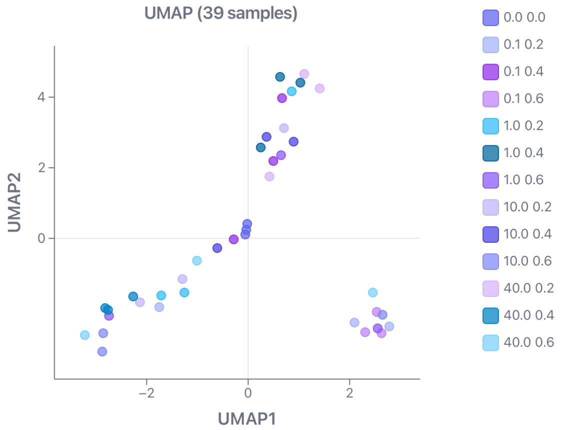


**B**

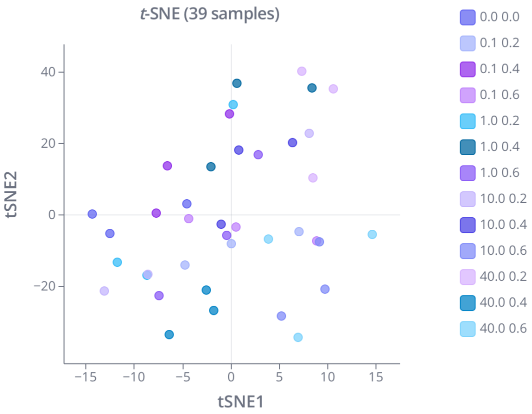


**Supplementary Fig. 7: UMAP & t-SNE of RNA Sequencing Groups.** UMAP (A) and t-SNE (B) plots display all samples in the dataset, including different mechanical index (MI)/microbubble volume dose (MVD) combinations and isoflurane control (0.0 0.0) samples.

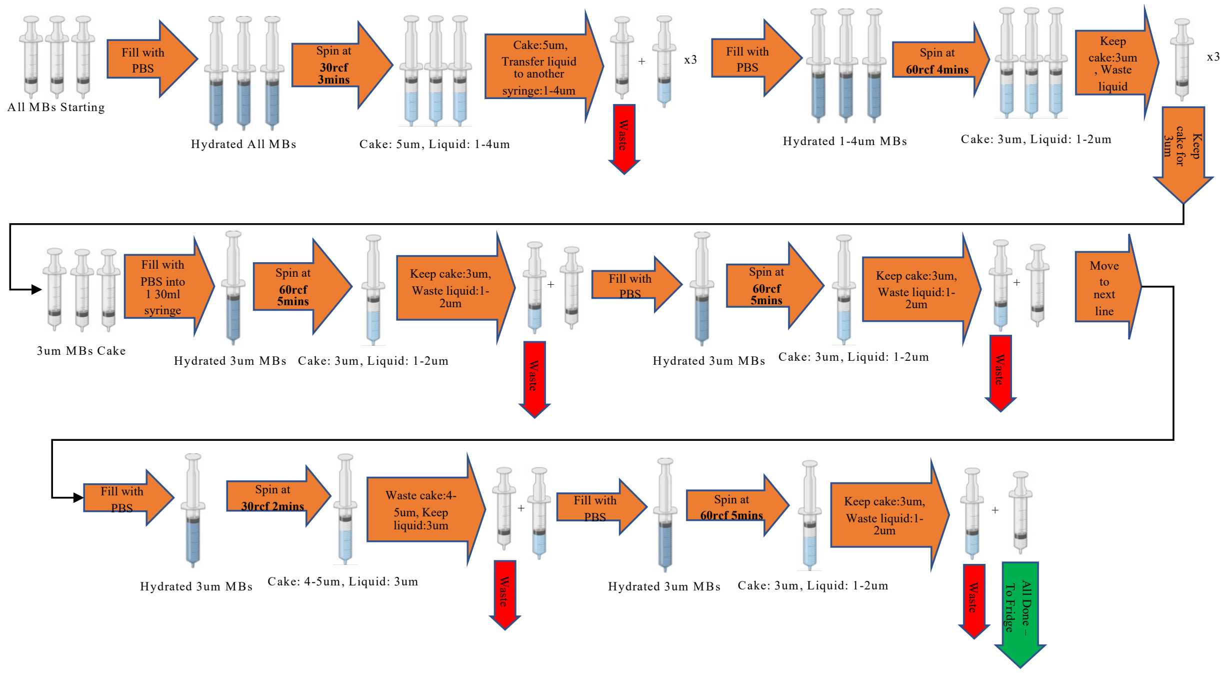


**Supplementary Fig. 8: Size Isolation Process of Microbubbles.** This is an illustration of the process used to isolate the 3 μm size distributions used in the study. The thicker orange arrow illustrates the separation of phases or centrifugation. Green arrows show when the cake was to be saved for measuring.

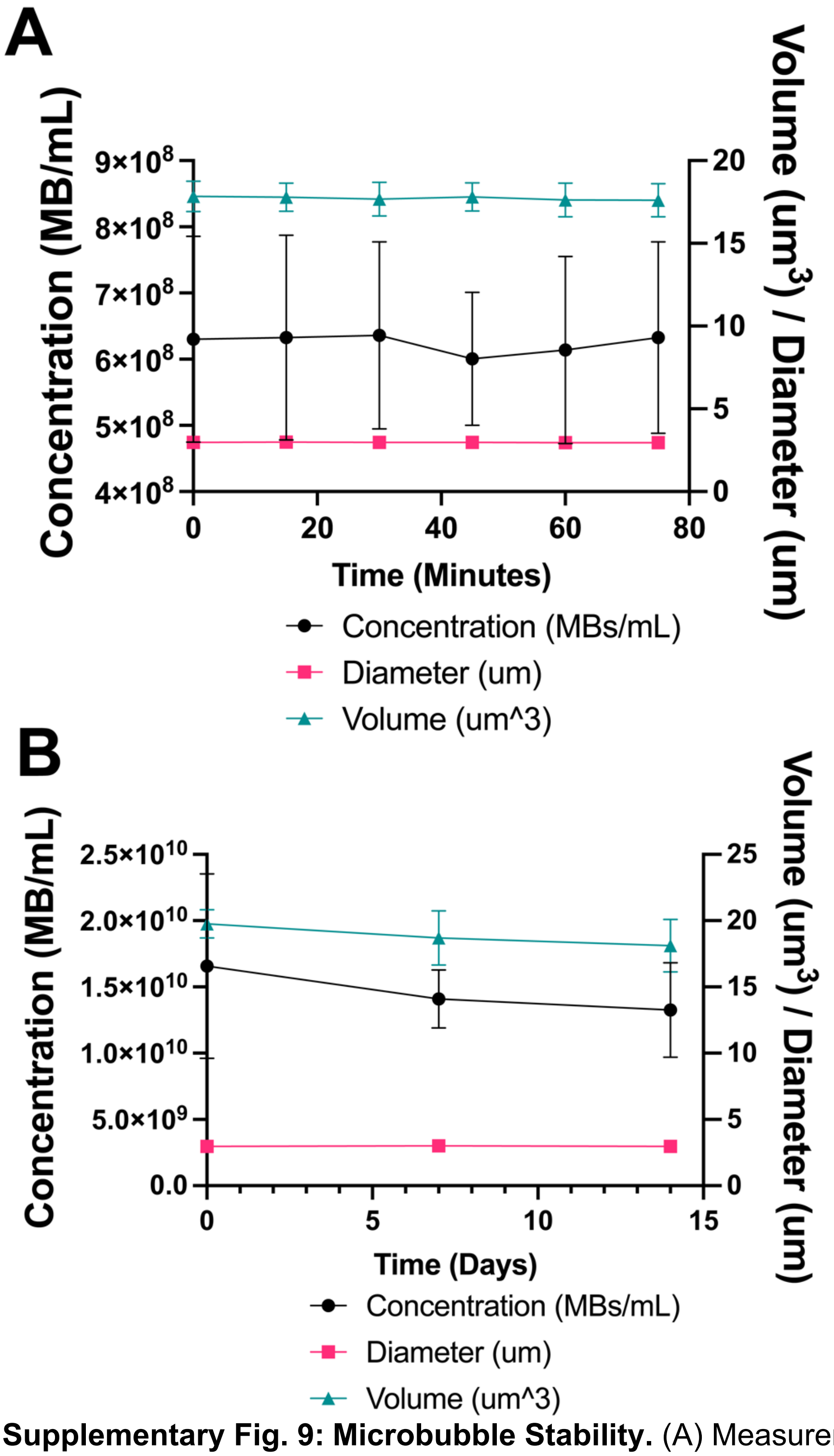


**Supplementary Fig. 9: Microbubble Stability.** (A) Measurements of microbubbles after 1 hour at diluted concentrations used during injection (~5E8MBs/mL). (B) The plot shows measurements of microbubbles after 2 weeks of storage at 4C. Data is shown as mean ± SD (*n = 3*).

### Supplementary Table 2: IHC Antibodies Used and Expected Cell Types.

| Antibodies | Species | Cell Type | Company | Catalog number | References |
|---|---|---|---|---|---|
| Iba1 | Rabbit | Microglia | CST | #17198 | (1, 2) |
| GFAP | Chicken | Astrocytes | Abcam | ab4674 | (1, 2) |
| CD4 | Rat | Helper T-Cells | Thermo | 42-0042-82 | (3, 4) |
| CD68 | Rat | Macrophages | Thermo | 14-0681-82 | (5, 6) |
| CD8 | Rabbit | Cytotoxic T-Cells | Thermo | MA514548 | (3, 7) |
| NFkB-p65 | Rabbit | N/A | Thermo | 51-0500 | (8) |
| CD44 | Rabbit | T-Cells | Thermo | 701406 | (2) |

### References.


1. S. Sinharay, *et al.*, In vivo imaging of sterile microglial activation in rat brain after disrupting the blood-brain barrier with pulsed focused ultrasound: [18F]DPA-714 PET study. *J Neuroinflammation* **16**, 155 (2019).

2. Z. I. Kovacs, *et al.*, Disrupting the blood–brain barrier by focused ultrasound induces sterile inflammation. *Proc. Natl. Acad. Sci. U.S.A.* **114** (2017).

3. Y.-Y. Fu, *et al.*, T Cell Recruitment to the Intestinal Stem Cell Compartment Drives Immune-Mediated Intestinal Damage after Allogeneic Transplantation. *Immunity* **51**, 90-103.e3 (2019).

4. S. U. Hridi, *et al.*, Increased Levels of IL-16 in the Central Nervous System during Neuroinflammation Are Associated with Infiltrating Immune Cells and Resident Glial Cells. *Biology* **10**, 472 (2021).

5. Z. I. Kovacs, *et al.*, MRI and histological evaluation of pulsed focused ultrasound and microbubbles treatment effects in the brain. *Theranostics* **8**, 4837–4855 (2018).

6. R. Waller, *et al.*, Iba-1-/CD68+ microglia are a prominent feature of age-associated deep subcortical white matter lesions. *PLoS ONE* **14**, e0210888 (2019).

7. S. Bhattacharya, K. Calar, C. Evans, M. Petrasko, P. De La Puente, Bioengineering the Oxygen-Deprived Tumor Microenvironment Within a Three-Dimensional Platform for Studying Tumor-Immune Interactions. *Front. Bioeng. Biotechnol.* **8**, 1040 (2020).

8. M. Liu, *et al.*, Isoflurane Conditioning Provides Protection against Subarachnoid Hemorrhage Induced Delayed Cerebral Ischemia through NF-kB Inhibition. *Biomedicines* **11**, 1163 (2023).
